# Expansion of profibrotic monocyte-derived alveolar macrophages in patients with persistent respiratory symptoms and radiographic abnormalities after COVID-19

**DOI:** 10.1101/2023.07.30.551145

**Authors:** Joseph I. Bailey, Connor H. Puritz, Karolina J. Senkow, Nikolay S. Markov, Estefani Diaz, Emmy Jonasson, Zhan Yu, Suchitra Swaminathan, Ziyan Lu, Samuel Fenske, Rogan A. Grant, Hiam Abdala-Valencia, Ruben J. Mylvaganam, Janet Miller, R. Ian Cumming, Robert M. Tighe, Kymberly M. Gowdy, Ravi Kalhan, Manu Jain, Ankit Bharat, Chitaru Kurihara, Ruben San Jose Estepar, Raul San Jose Estepar, George R. Washko, Ali Shilatifard, Jacob I. Sznajder, Karen M. Ridge, GR Scott Budinger, Rosemary Braun, Alexander V. Misharin, Marc A. Sala

## Abstract

As many as 10–30% of the over 760 million survivors of COVID-19 develop persistent symptoms, of which respiratory symptoms are among the most common. To understand the cellular and molecular basis for respiratory PASC, we combined a machine learning based analysis of lung computed tomography (CT) with flow cytometry, single-cell RNA-sequencing analysis of bronchoalveolar lavage fluid and nasal curettage samples, and alveolar cytokine profiling in a cohort of thirty-five patients with respiratory symptoms and radiographic abnormalities more than 90 days after infection with COVID-19. CT images from patients with PASC revealed abnormalities involving 73% of the lung, which improved on subsequent imaging. Interstitial abnormalities suggestive of fibrosis on CT were associated with the increased numbers of neutrophils and presence of profibrotic monocyte-derived alveolar macrophages in BAL fluid, reflecting unresolved epithelial injury. Persistent infection with SARS-CoV-2 was identified in six patients and secondary bacterial or viral infections in two others. These findings suggest that despite its heterogenous clinical presentations, respiratory PASC with radiographic abnormalities results from a common pathobiology characterized by the ongoing recruitment of neutrophils and profibrotic monocyte-derived alveolar macrophages driving lung fibrosis with implications for diagnosis and therapy.

## INTRODUCTION

Over 768 million patients have been infected with SARS-CoV-2, and over 6.9 million people have died from their infection since 2020^1^. Despite the availability of effective vaccines and widespread immunity in the population, SARS-CoV-2 remains an important cause of viral pneumonia and the emergence of new SARS-CoV-2 variants or other coronavirus strains remains a public health concern^2^. Many patients who had COVID-19 continue to have symptoms months after their initial infection^3^. Termed post-acute sequelae of COVID-19 (PASC), these patients may suffer from a wide variety of symptoms across organ systems^4^. A subset of patients with PASC present with shortness of breath, cough, or hypoxemia and detectable abnormalities on computed tomography (CT) scan^5^, which we refer to as respiratory PASC with radiographic abnormalities (RPRA). Current practice guidelines suggest clinicians consider treating these patients with systemic corticosteroids, but these guidelines are based on weak evidence and the molecular underpinnings of RPRA are largely unexplored^6, 7^. Furthermore, CT findings in patients with RPRA are heterogenous both within and between patients and include areas of parenchymal fibrosis, ground glass opacities, or organizing pneumonia, among others^8^. These findings leave open the question of whether RPRA represents heterogenous clinical phenotypes with a shared pathobiology or a syndromic collection of disorders with distinct pathobiologies.

Previously, we applied a systems biology approach to analyze bronchoalveolar lavage (BAL) samples from patients with severe acute COVID-19. We discovered self-amplifying inflammatory circuits between alveolar macrophages harboring the virus and activated T cells^9^. Our group and others have causally linked these monocyte-derived alveolar macrophages to the development of lung injury and fibrosis in animal models^10–14^. We confirmed the presence of a homologous population of alveolar macrophages in the lungs of patients with pulmonary fibrosis who required lung transplantation, including patients with post-COVID-19 pulmonary fibrosis^11, 15^. Furthermore, using mouse models of pulmonary fibrosis, we and others have found the recruitment of monocyte-derived alveolar macrophages is interrupted by successful alveolar repair^16, 17^. Based on these findings, we hypothesized that RPRA results from failed alveolar epithelial repair after SARS-CoV-2 pneumonia that can be detected by the ongoing recruitment of profibrotic monocyte-derived alveolar macrophages in BAL samples. If so, these cells might provide a useful biomarker and a potential therapeutic target.

Accordingly, we conducted an observational study that included 35 patients with RPRA in which we analyzed serial CT images using a validated machine learning approach^18, 19^ and combined these data with flow cytometry, single-cell RNA-seq, and cytokine analysis of BAL fluid. We found evidence of significant radiographic lung abnormalities at the time of presentation. Flow cytometry and single-cell RNA-seq analysis showed the ongoing recruitment of neutrophils and profibrotic monocyte-derived alveolar macrophages associated with the degree of fibrotic abnormalities on CT scans. In six patients, SARS-CoV-2 was detected by PCR in BAL fluid and in three of these patients viral transcripts were detected using single-cell RNA-seq. Serial CT imaging showed lung abnormalities improved in most patients in response to treatment and time, with the largest improvement observed in areas suggestive of fibrosis. Our findings suggest a failure to resolve SARS-CoV-2-induced alveolar inflammation contributes to RPRA in some patients and supports microbiologic and molecular evaluation of the alveolar space as potential diagnostic tools to inform the treatment of these patients.

## RESULTS

### Study cohort

The study cohort included 35 patients with respiratory PASC with radiographic abnormalities (RPRA), including two patients with RPRA and severe lung fibrosis who ultimately required lung transplantation. Bronchoalveolar lavage (BAL) fluid samples from two separate cohorts of 12 and 9 healthy volunteers and nasal mucosal curettage from 6 healthy volunteers were used as a comparison. Demographic features of the patients and volunteers are summarized in **Supplemental Table 1** and **Supplemental Data Files 1 and 2**. The median age of the patients with RPRA was 62 (range from 32 to 83), 43% of the patients were female, 66% of the patients were white, 17% black or African American and 20% self-identified as Hispanic or Latino. The clinical course of the patients in the cohort noting treatments is shown in **Figure 1a**. All patients had respiratory symptoms on presentation with shortness of breath being the most common symptom (97%) followed by cough (69%). Fifteen patients (43%) exhibited arterial hypoxemia requiring oxygen therapy, including both of the patients who required lung transplantation (**Table 1**). Nine patients had never been hospitalized, of those hospitalized, 17 were admitted to the ICU during their hospitalization. A bronchoscopy was performed on 30 patients, and the median time from the diagnosis of COVID-19 to bronchoscopy was 201±74 days (**Figure 1a,b**). Twenty-seven patients received steroids for the treatment of RPRA. Of those who received corticosteroids and underwent bronchoscopy, 18 underwent a bronchoscopy before treatment, and three were undergoing or had completed treatment before the bronchoscopy. Five patients were treated with steroids without undergoing bronchoscopy (**Figure 1a,b**).

**Figure 1:**
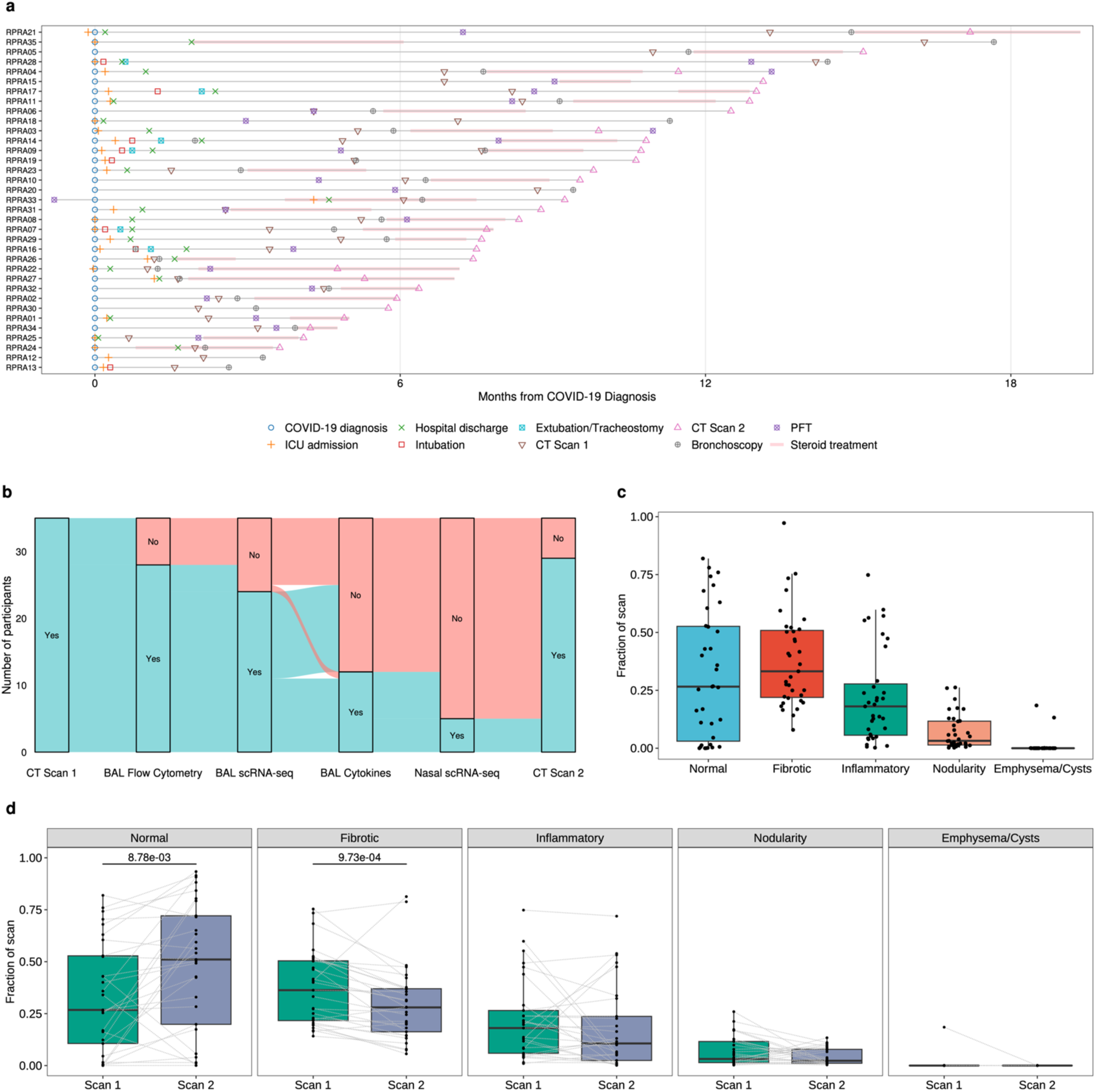
Patients with RPRA exhibit fibrotic abnormalities on CT imaging that improve with time. **a.** Summary of the clinical course of the cohort. Key events are annotated as indicated in the legend. **b.** Sankey diagram illustrating analyses performed for the cohort. **c.** Computed tomography (CT) scan abnormalities were quantified using a machine learning algorithm and the reported abnormalities were grouped into normal lung, fibrotic abnormalities, inflammatory abnormalities, nodularity, and emphysema or cysts (see Table S3). **d.** Changes in abundance of radiographic abnormalities over time. Scans of the same subject are connected. Each CT scan is represented by a single point. Adjusted p-values are shown above pairs of boxplots when changes were significant (q < 0.05, paired Wilcoxon rank-sum tests with FDR correction).

**Table 1.**
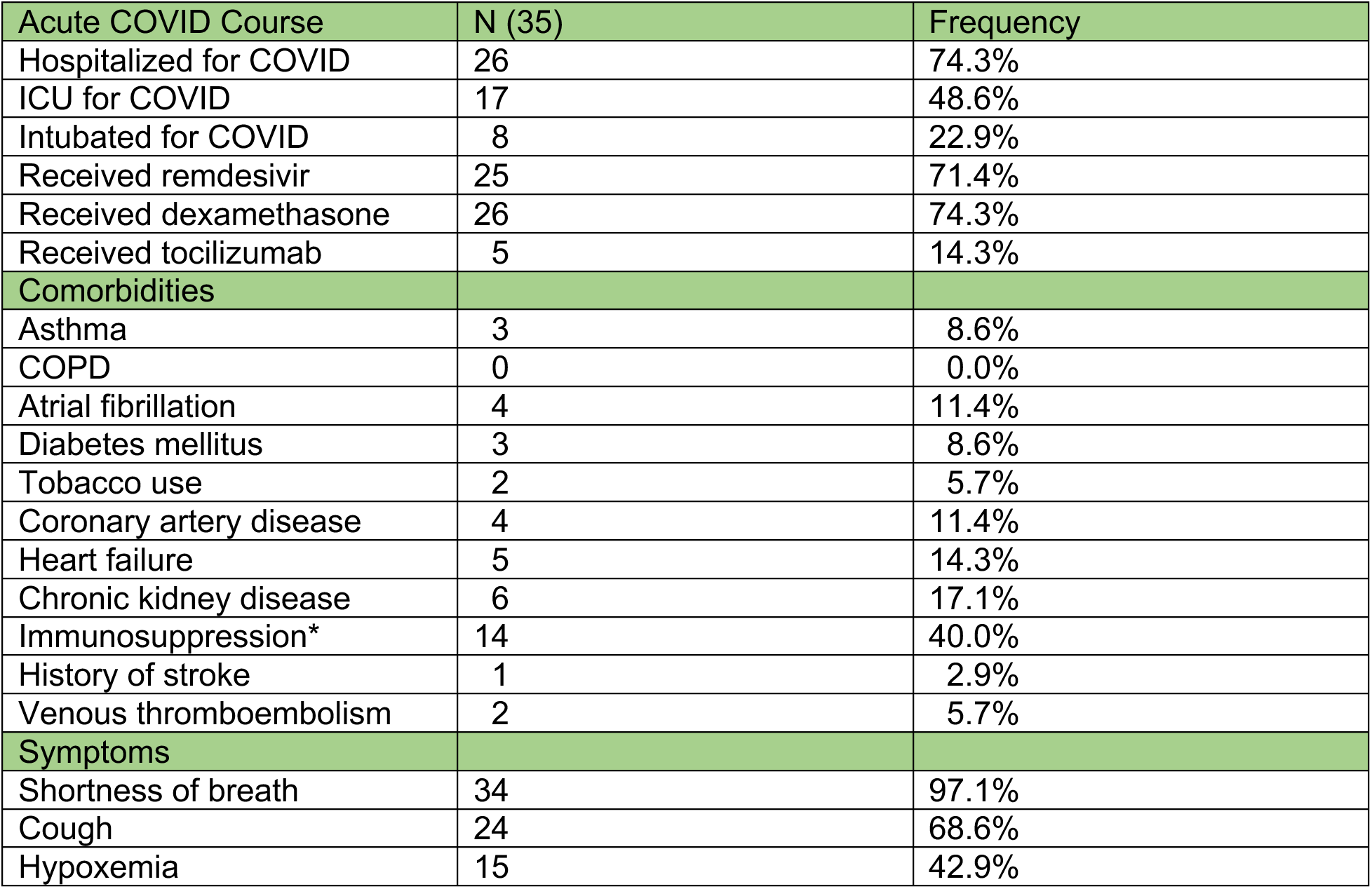
Clinical features of the cohort. Immunosuppression was defined as receiving medications known to cause immunosuppression at time of RPRA evaluation, such as glucocorticoids, mycophenolate, cyclosporine, or tacrolimus.

| Acute COVID Course | N (35) | Frequency |
| --- | --- | --- |
| Hospitalized for COVID | 26 | 74.3% |
| ICU for COVID | 17 | 48.6% |
| Intubated for COVID | 8 | 22.9% |
| Received remdesivir | 25 | 71.4% |
| Received dexamethasone | 26 | 74.3% |
| Received tocilizumab | 5 | 14.3% |
| **Comorbidities** |  |  |
| Asthma | 3 | 8.6% |
| COPD | 0 | 0.0% |
| Atrial fibrillation | 4 | 11.4% |
| Diabetes mellitus | 3 | 8.6% |
| Tobacco use | 2 | 5.7% |
| Coronary artery disease | 4 | 11.4% |
| Heart failure | 5 | 14.3% |
| Chronic kidney disease | 6 | 17.1% |
| Immunosuppression* | 14 | 40.0% |
| History of stroke | 1 | 2.9% |
| Venous thromboembolism | 2 | 5.7% |
| **Symptoms** |  |  |
| Shortness of breath | 34 | 97.1% |
| Cough | 24 | 68.6% |
| Hypoxemia | 15 | 42.9% |

### Computed tomography of the chest reveals extensive abnormalities in patients with RPRA

All patients included in this study had abnormalities visible using computed tomography (CT) of the chest. Radiographic abnormalities extracted from the official radiology interpretations of the initial CTs are included in **Supplemental Table 2**. Because standard clinical interpretation of CT is qualitative, we used a previously established machine learning algorithm^18, 19^ to quantify the radiographic abnormalities seen on the initial CT scans (**Supplemental Data File 3**). This procedure quantifies abnormal regions of the lung using a machine learning classifier and normalizes the abnormalities to the estimated lung volume. We grouped these abnormalities into those recognized by clinicians as areas characterized by normal lung, lung fibrosis, lung inflammation, emphysema/cysts or lung nodularity (**Supplemental Table 3** and **Supplemental Data Files 4-5**).

Quantitative assessment of the initial CT scans revealed significant abnormalities, with only 31.6±27.6% of the lung classified as “normal” (**Figure 1c**). The most commonly observed abnormalities were those typically thought to represent changes compatible with fibrosis (38.2±20.4%), followed by increased parenchymal attenuation compatible with inflammation (22.3±20.3%) and nodularity (7.0±7.5%) (**Figure 1c**)^20^. Two patients had abnormalities classified as emphysema on the first scan. One of these patients did not have a second scan. The second patient had findings of emphysema on the first scan, which was not present on the subsequent scan. Manual review of the initial scan suggested hyperinflation rather than tissue destruction underlying the reduced density.

Twenty-nine of the patients underwent a second CT scan. The median time between the first and second CT scan was 118±51.7 days. Analysis of the second compared with the first CT scan showed significant improvement (normal area increased to 48.1±31.5%) (**Figure 1d**). The bulk of the improvement was attributable to reduced fibrotic abnormalities. There was no association with the amount of improvement (measured as change in the percentage of normal lung) and the interval between the CT scans (**Supplemental Figure 1**).

### RPRA is associated with an increased relative abundance of neutrophils and monocytes in bronchoalveolar lavage fluid

We used flow cytometry to compare the cellular composition of BAL fluid in 9 healthy volunteers and 28 of the patients with RPRA (**Figure 2a**). We identified neutrophils, eosinophils, CD4+ and CD8+ T cells, regulatory T cells (Tregs), monocytes and macrophages in the BAL fluid. Since newly differentiated monocyte-derived alveolar macrophages express lower levels of mannose receptor CD206^9^, we separated alveolar macrophages into more mature CD206^high^ and less mature CD206^low^ alveolar macrophages. Hierarchical clustering on cell type abundance identified four clusters of samples (**Figure 2a, Supplemental Data File 6**). Cluster 1 was characterized by an increased abundance of neutrophils, monocytes, and CD8 T cells. Both patients who underwent lung transplant for post-COVID-19 fibrosis were in Cluster 1. Cluster 2 was dominated by CD206^high^ alveolar macrophages and was composed primarily of samples from healthy volunteers. Clusters 3 and 4 had intermediate phenotypes: Cluster 3 had an increased abundance of CD206^low^ alveolar macrophages and eosinophils, while Cluster 4 was characterized by an increased abundance of Tregs, CD4 T cells and NK cells. Clusters 1 and 3 were composed entirely of patients with RPRA, while more than half of Cluster 4 was patients with RPRA. Direct pairwise comparison between healthy volunteers and patients with RPRA demonstrated a significant increase in the relative abundance of neutrophils and monocytes, and a decrease in the relative abundance of CD206^high^ and CD206^low^ alveolar macrophages (**Figure 2b; Supplemental Figure 2a**).

**Figure 2:**
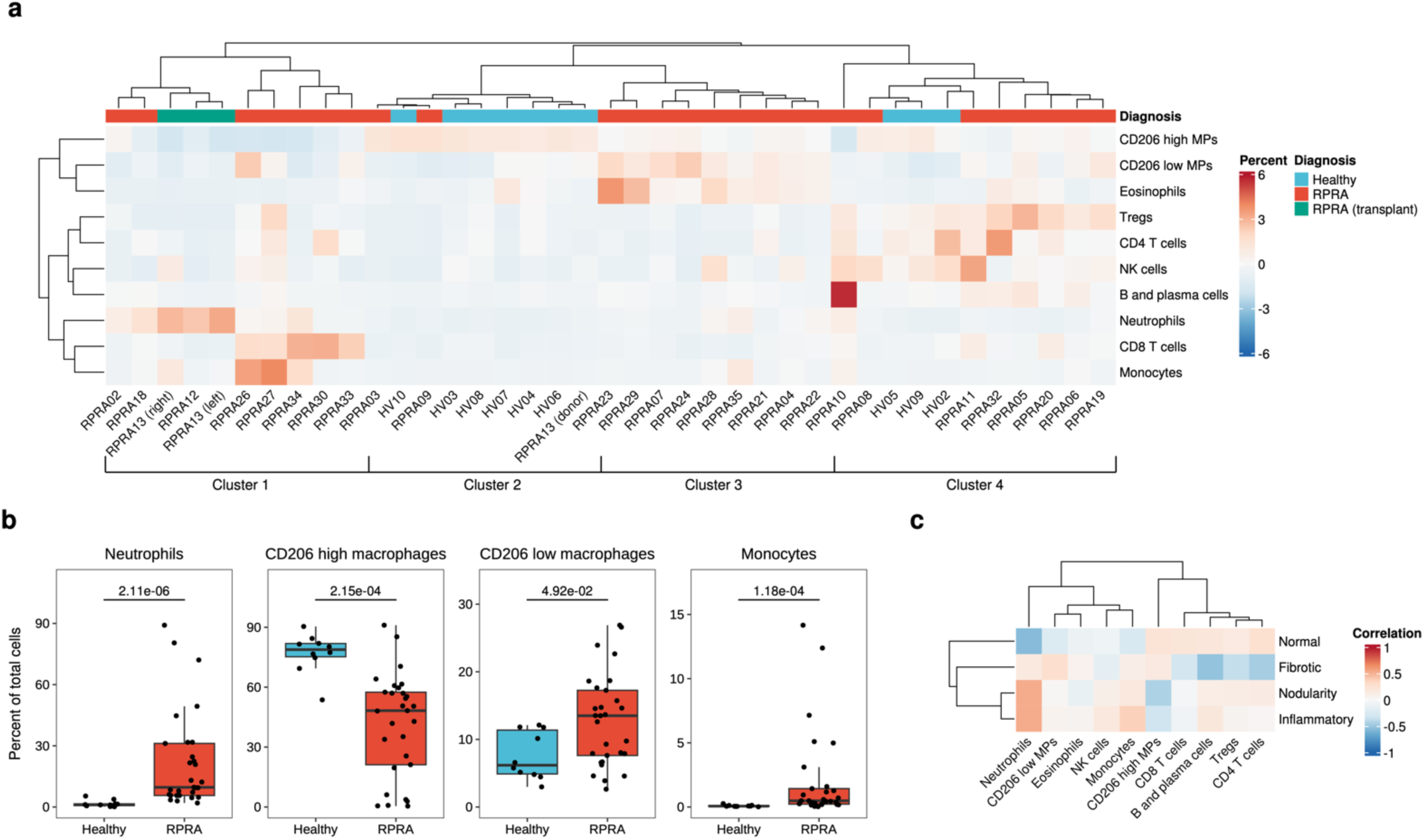
Flow cytometry shows an expansion of monocyte-derived alveolar macrophages and neutrophils in patients with RPRA compared with healthy volunteers. **a.** Hierarchical clustering of flow cytometry data from BAL samples from healthy volunteers and patients with RPRA. Column headers are color-coded by the diagnosis. Samples from the two patients with RPRA who required lung transplant are coded separately (one of these patients had two BAL samples, where BAL fluid was obtained from each lung separately). Clustering was performed using Ward’s method. Rows are z-scored. MP, macrophage. **b.** Proportions of significantly differentially abundant cells in BAL fluid between healthy controls and patients with RPRA (including patients who required lung transplant) (q < 0.05, pairwise Wilcoxon rank-sum tests with FDR correction). Adjusted p-values are shown above each pair of boxplots. **c.** Hierarchical clustering of correlation coefficients (Spearman’s rho) between cell type abundances and fraction of CT features. Clustering was performed using Ward’s method. No associations were statistically significant (q < 0.05, permutation tests with FDR correction).

We then evaluated the correlation between the fraction of radiographic features extracted from CT images and the abundance of immune populations in BAL fluid in patients with RPRA (**Figure 2c**). The area of normal lung on the initial CT scan inversely correlated with abundance of neutrophils, while the area of inflammatory and nodular abnormalities positively correlated with abundance of neutrophils. In contrast, relative abundance of regions of nodular abnormalities was negatively associated with an abundance of CD206^high^ alveolar macrophages, and relative abundance of fibrotic abnormalities was negatively associated with abundance of CD4 T cells, Tregs, and B and plasma cells (**Figure 2c, Supplemental Figure 2b**). However, these correlations were not statistically significant.

### Single-cell RNA-seq of immune cells from bronchoalveolar lavage fluid shows ongoing recruitment of profibrotic monocyte-derived alveolar macrophages associated with radiographic severity in patients with RPRA

We have previously reported that monocyte-derived alveolar macrophages (MoAM) play a causal role in the development of lung fibrosis in mouse models^10, 12, 16^. We identified a homologous population in lung explants and autopsy specimens from patients with end-stage idiopathic pulmonary fibrosis and end-stage post-COVID-19 pulmonary fibrosis^11, 15^. We therefore hypothesized that profibrotic MoAMs would be present in BAL fluid from patients with RPRA and would reflect disease severity. Accordingly, we performed single-cell RNA-seq on flow-sorted mononuclear immune cells from BAL fluid obtained from 24 patients in our cohort – including the patients who required lung transplant – and 6 healthy volunteers. Integrative analysis resolved all cell types previously reported in BAL fluid: macrophages, subsets of dendritic cells, subsets of T cells, mast cells, B cells and plasma cells, as well as a cluster of SARS-CoV-2 particles (**Figure 3a,b, Supplemental Figure 3a–b, Supplemental Data File 7,** single-cell RNA-seq data can be explored at https://www.nupulmonary.org/)^21,22^. We found that after adjustment for the proportion of neutrophils (which were excluded from single-cell RNA-seq analysis) the cell type abundance determined by flow cytometry and single-cell RNA-seq correlated well with each other (**Supplemental Figure 3c**). Within the macrophage cluster (expressing *C1QA, MRC1, MSR1*), we resolved mature tissue-resident alveolar macrophages (TRAM), characterized by expression of *FABP4* and *INHBA;* and MoAMs, characterized expression of *VCAN* and *CCL2* and lack of expression of *FABP4* (**Figure 3c, Supplemental Figure 3d**)^21, 23, 24^. Interestingly, while these samples were collected from alveolar space via BAL, we detected a cluster of macrophages characterized by the expression of *SELENOP, CCL13, FOLR2*, which matched gene expression profiles of perivascular macrophages^21, 25^.

**Figure 3:**
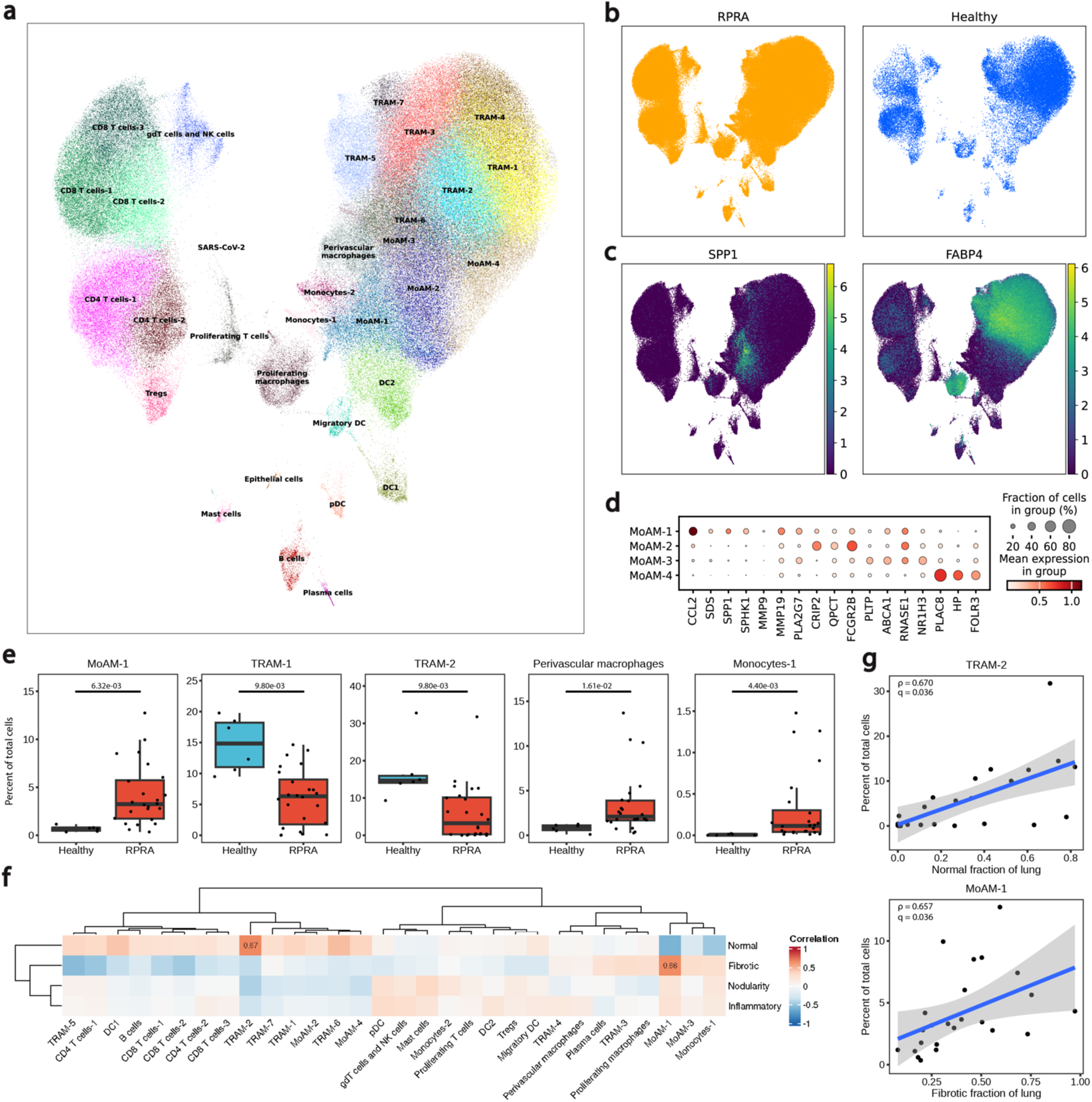
Ongoing recruitment of profibrotic MoAMs is associated with fibrotic abnormalities on CT scans. **a.** UMAP plot showing integrated analysis of BAL immune cells from 24 patients with RPRA and 6 healthy volunteers. **b.** UMAP plots of cells only in RPRA patients, and only in healthy volunteers. **c.** Expression of *SPP1* and *FABP4*. **d.** Dot plot showing expression of marker genes distinguishing subsets of MoAMs. **e.** Proportions of significantly differentially abundant cells in BAL fluid between healthy controls and patients with RPRA (including patients who required lung transplant) (q < 0.05, pairwise Wilcoxon rank-sum tests with FDR correction). Adjusted p-values are shown above each pair of boxplots. **f**. Hierarchical clustering on correlation coefficients (Spearman’s rho) between cell type abundances determined using single-cell RNA-seq and CT features. Correlation coefficients are shown only when the association was significant (q < 0.05, permutation tests with FDR correction). Clustering was performed using Ward’s method. **g**. Correlation between cell type abundance for the described cell populations with the described CT features. Only significant associations are shown, with adjusted p-values and correlations shown on each plot. Linear models and 95% confidence intervals are shown.

We resolved two subsets of monocytes (Monocytes-1 *CD300E, CCL2, FCN1, VCAN* and Monocytes-2 *APOBEC3A, IFITM3, CXCL11*) as well as four subsets of MoAMs (MoAM-1–4). MoAM-1 were the least mature and were characterized by expression of *FCN1, VCAN, CCL2, MSR1* and low expression of *C1QA*. In addition, MoAM-1 were characterized by high expression of the genes which have been detected in macrophages isolated from patients with pulmonary fibrosis^11, 24, 26–28^ and have been causally implicated in the pathogenesis of pulmonary fibrosis in animal and *in vitro* models, such as *SPP1, SPHK1, PLA2G7, MMP19* and others (**Figure 3c,d, Supplemental Figure 3d**)^28–33^. We thus refer to this cluster as profibrotic MoAM-1. Clusters MoAM-2–4 showed progressively increasing expression of genes associated with macrophage maturation and adaptation to alveolar niche (**Figure 3d**).

To further characterize clusters of immune cells identified in BAL, we performed factor analysis using the Spectra method^34^ to decompose the single-cell RNA-seq count matrix into a set of interpretable gene programs or factors. Taking a curated list of factors as input, Spectra both modifies them and builds novel factors to help explain the observed variation in expression. We provided Spectra with 68 factors related to cellular identity and cellular processes in both generic immune cells and in specific cell populations. Additionally, we allowed Spectra to discover one novel factor for each broad cell population for a total of 77 factors (**Supplemental Figure 3e**, **Supplemental Data Files 8-9**). Profibrotic MoAM-1 were characterized by the expression of partially overlapping programs F0, F9, and F17, which in addition to already mentioned profibrotic genes (such as *SPP1, CHI3L1, MMP9*) included genes related to mitochondrial respiratory chain function (*C15orf48*), antioxidant activity (*SOD2*), iron metabolism and homeostasis (*HAMP*), immune response and inflammation (*IL1RN, FCN1, CD83, LYZ*), lipid metabolism and transport (*APOE, APOC1, APOC2, FABP5*), amino acid metabolism (*SDS*), extracellular matrix remodeling and degradation (*CTSL*, *VCAN, LGMN, MARCO*), cell adhesion and cytoskeletal organization (*MARCKS, EZR, EMP1*), and cell signaling and migration (*FCGR2B*, *SGK1, NR4A3, CD48, CD14, SPINK1, PHLDA1, IFITM3, IFI30, CXCR4, MS4A6A*).

We compared the differential abundance of cells within each single-cell RNA-seq cluster in patients with RPRA (including patients with COVID-19-induced lung fibrosis who required lung transplantation) and healthy controls. We found that the relative abundance of Monocytes-1, MoAM-1, and perivascular macrophages was significantly increased while the relative abundance of TRAM-1 and TRAM-2 was significantly decreased (**Figure 3e, Supplemental Figure 4a,b, Supplemental Data File 11**). We then examined the relationship between the abundance of different subclusters identified from our single-cell RNA-seq analysis and the fraction of abnormal lung on CT scan. The abundance of profibrotic MoAM-1 negatively correlated with area of normal lung on CT scans and positively correlated with fibrotic lung abnormalities on CT scan, while the abundance of TRAM-2 positively correlated with area of the normal lung (**Figure 3f,g)**.

### Transcriptomic changes in alveolar macrophages reflect radiographic abnormalities in RPRA

While profibrotic MoAM-1 were present nearly exclusively in patients with RPRA and almost absent in healthy volunteers, thus precluding differential gene expression analysis, TRAMs were present in both groups. Furthermore, the abundance of TRAMs subsets detected using either flow cytometry or single-cell RNA-seq was positively correlated with the fraction of normal lung on CT. As these cells are thought to be important sensors of changes in alveolar homeostasis, we compared gene expression within TRAM populations in patients with RPRA and healthy controls. Very few differentially expressed genes were shared across TRAM subsets (**Figure 4a, Supplemental Figure 4c,d, Supplemental Data File 12**). We focused on TRAM-1 and TRAM-2 populations, which were differentially abundant between healthy volunteers and patients with RPRA. Among 44 genes upregulated in TRAMs from patients with RPRA were genes encoding components of the mitochondrial electron transport chain (*NDUFA7, NDUFB7, UQCRB, NDUFC1, NDUFB4, NDUFS6, COX7C, COX6A1, COX6B1, COX5B, COX8A*) or mitochondria organization (*MICOS13*), alarmins (*S100A6, S100A11*), and response to interferon (*IFNGR2, IFI27L2*). Among 12 downregulated genes were “classical” homeostatic TRAM genes, including *CSF2RB* (which encodes subunit of CSF2 receptor, a key cytokine for alveolar macrophage differentiation and survival), *TFRC* (transferrin receptor), genes involved in lipid and glucose metabolism (*ESYT1, PYGL, HK2*), and DNA repair and modification (*PARP10, RFC2*) (**Figure 4b**).

**Figure 4:**
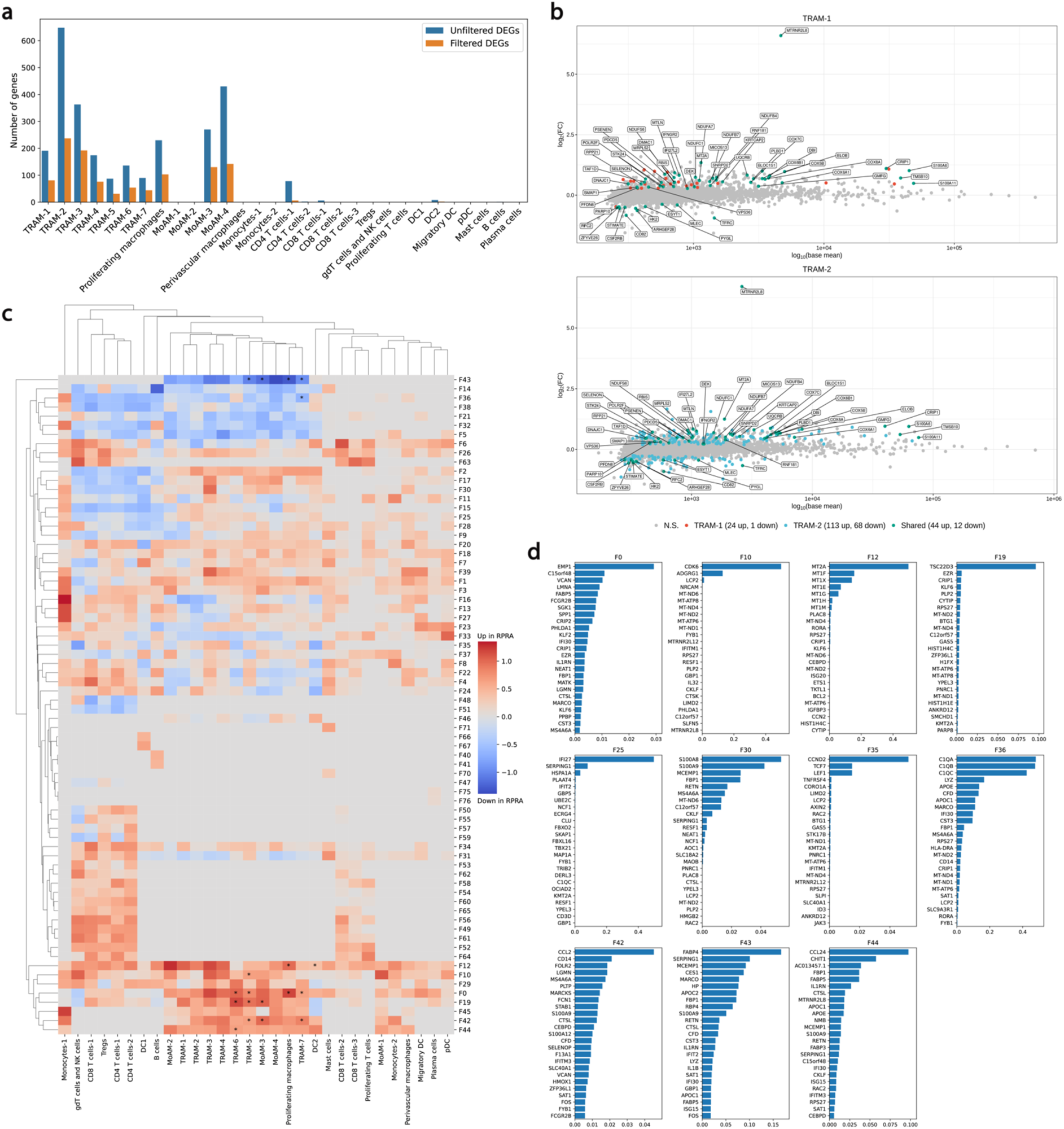
Differential gene expression in BAL macrophages in patients with RPRA. **a.** Bar plot showing number of differentially expressed genes between different cell types in BAL fluid in healthy controls and patients with RPRA (q < 0.05, Wald test with FDR correction) with and without filtering criteria applied (see methods). **b.** MA plots of all differentially expressed genes in TRAM-1 and TRAM-2 populations. Genes upregulated or downregulated in both populations are annotated. N.S., not significant (q > 0.05). **c.** Hierarchical clustering on the signal-to-noise ratio of Spectra subject scores between patients with RPRA and healthy controls. Factors which are differentially expressed (q < 0.05, Wilcoxon rank-sum test on subject scores with FDR correction) are indicated with an asterisk. **d.** Gene weights of the top 25 genes by weight of selected factors.

We compared expression of gene programs identified by Spectra algorithm^34^ between healthy volunteers and patients with RPRA and found 7 differentially expressed programs (**Figure 4c**). Program F43, containing genes related to homeostatic alveolar macrophage functions was downregulated in MoAM-3, TRAM-5, TRAM-7, and proliferating macrophages from patients with RPRA compared with healthy controls. Three partially overlapping programs, F0, F19, and F42 were upregulated in TRAM-5 and TRAM-6 from patients with RPRA compared with healthy controls. These included genes involved in immune responses, cell adhesion and cytoskeletal organization, extracellular matrix organization and fibrosis, protease activity, RNA regulation, DNA and chromatin regulation, and iron metabolism (**Figure 4d, Supplemental Data File 10**).

We then asked whether expression level of gene programs identified by Spectra correlated with degree and type of radiographic abnormalities in patients with RPRA (**Supplemental Figure 4e**). Interestingly, we found that expression of program F35 in Tregs negatively correlated with the amount of normal appearing lung parenchyma on CT. This program included genes related to various processes such as cell cycle regulation and proliferation, Wnt-signaling, immune response, cell adhesion and migration and signaling. As Tregs are causally linked to lung repair in animal models^35, 36^, these findings might suggest dysfunction in these cells contributes to failed resolution^37, 38^. Expression of program F30 in TRAM-6, related to inflammation, immune response, pathogen clearance, metabolic regulation, and complement regulation, was positively correlated with the fraction of normal lung. Expression of program F25 in MoAM-3 and MoAM-4, associated with immune response, antiviral defense, complement regulation, lysosomal activity and degradation negatively correlated with the fraction of fibrotic radiographic abnormalities. Combined these findings suggest activation of macrophage and T cell populations are associated with the detection of inflammatory and fibrotic abnormalities in patients with RPRA.

### Cytokine profiles in BAL fluid identify persistent lung inflammation in patients with RPRA

We measured the levels of 71 cytokines and chemokines in BAL fluid from 12 patients with RPRA and 13 healthy controls using a multiplex bead array. Of these, 43 were detected in the BAL fluid samples and were included in the subsequent analysis (see Methods). Compared to healthy controls, the levels of 13 cytokines were significantly higher and three were significantly lower in BAL fluid from patients with RPRA (**Figure 5a,b, Supplemental Figure 5a, Supplemental Data File 13**). Upregulated molecules included monocyte and T cell chemoattractants MCP-1 (*CCL2*), MCP-3 (*CCL7*), MIP-1β (*CCL4*), MIP-1δ (*CCL15*) and the inflammatory cytokines IL-1ɑ, IL-1RA, IL-6 and IL-8. Also upregulated was BCA-1 (*CXCL13*) involved in B lymphocyte recruitment; EGF which promotes epithelial repair; ENA-78 (*CXCL5*) implicated in neutrophil recruitment during repair; eotaxin-1 (*CCL11*) involved in eosinophil recruitment; a member of the TNF superfamily soluble CD40 ligand (*CD40L*), which has broad effect on activation of many immune cells; and the apoptosis ligand TRAIL. Downregulated molecules included IL-13, involved in TH2-mediated inflammation; and IL-21, a cytokine involved in monocyte differentiation and T and NK cell activation. The upregulation of monocyte, T cell and neutrophil chemoattractants in RPRA supports the role of the corresponding cells as biomarkers or drivers of RPRA.

**Figure 5:**
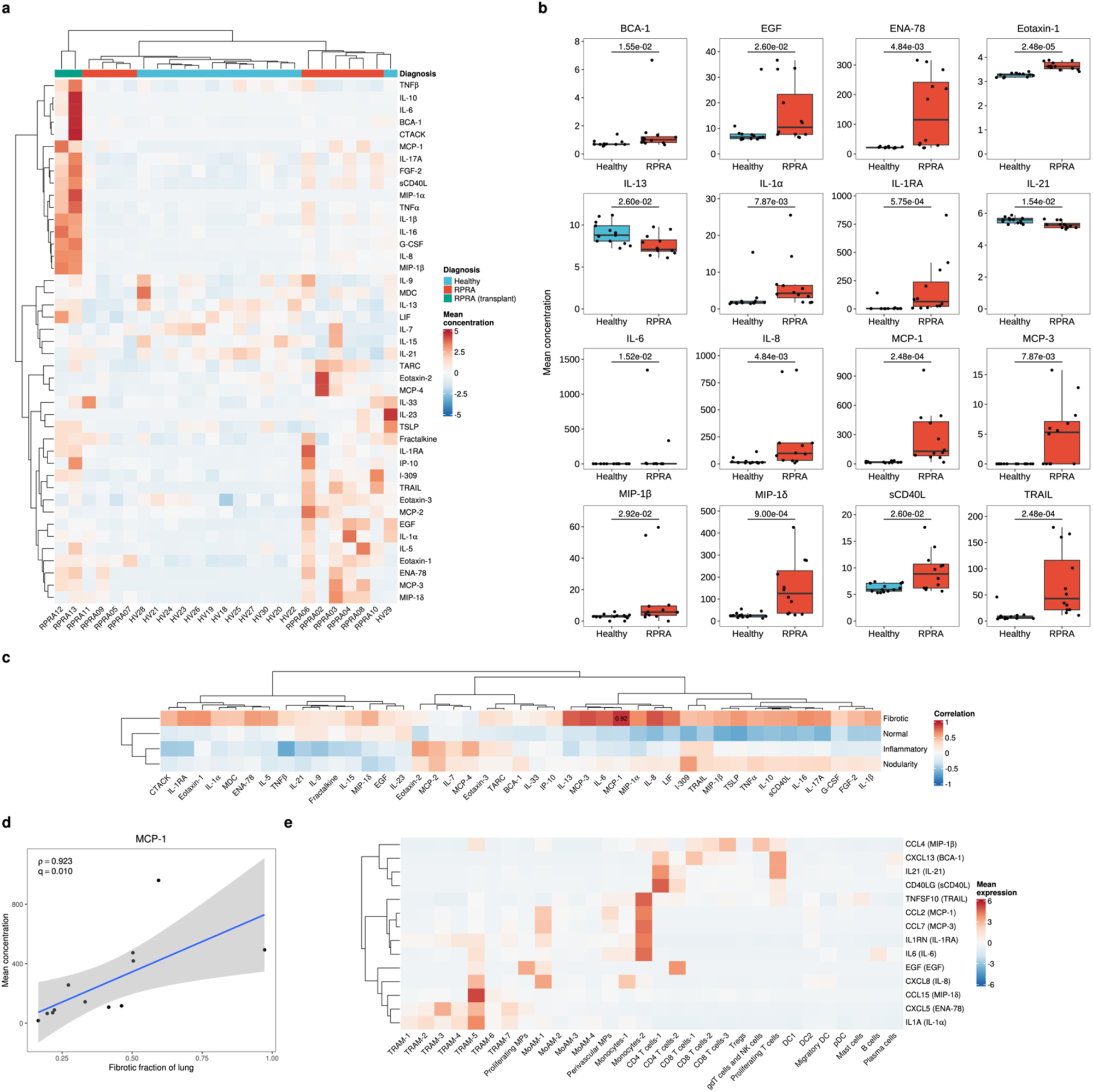
BAL fluid cytokines implicate monocyte and neutrophil cytokines and chemokines in RPRA. **a.** Hierarchical clustering of BAL fluid cytokines in patients with RPRA compared with patients with healthy controls. Patients with severe RPRA who required transplant are indicated as RPRA (transplant). A total of 71 cytokines were measured, results are shown for the 43 that were detected in any sample. Clustering was performed using Ward’s method. Rows were z-scored. **b.** Levels of 16 cytokines or chemokines that differed significantly (q < 0.05, pairwise Wilcoxon rank-sum tests with FDR correction) between patients with RPRA (including patients who required lung transplant) and healthy controls. Adjusted p-values are shown above each pair of boxplots. **c**. Hierarchical clustering on correlation coefficients (Spearman’s rho) between levels of inflammatory cytokines and CT features. Clustering was performed using Ward’s method. Correlation coefficients are shown only when the association was significant (q < 0.05, permutation tests with FDR correction). **d**. Comparison of cytokine levels with the described CT features. Only significant associations are shown, with correlations and adjusted p-values shown. A linear model and 95% confidence interval are shown. **e**. Hierarchical clustering of mean expression levels from BAL single-cell RNA-seq data of genes encoding each cytokine that differed significantly between patients with RPRA and healthy controls. *CCL11* (eotaxin-1) is not shown as it was not expressed in cells sampled via BAL procedure. Clustering was performed using Ward’s method. Rows are z-scored.

We then compared the levels of inflammatory cytokines within the group of patients with RPRA to the levels of CT abnormalities. While the levels of many of the cytokines were positively correlated with the severity of fibrosis measured on CT, this correlation was only significant for MCP-1 (*CCL2*), a chemokine central to the recruitment of monocytes (**Figure 5c,d**). We next aimed to identify potential source of the inflammatory cytokines that were elevated in the BAL of patients with RPRA compared with healthy controls. Many of the cytokines we found were elevated in bronchoalveolar lavage fluid from patients with RPRA were expressed in immune cells sampled by the BAL procedure (**Figure 5e, Supplemental Figure 5b**). For example, MoAMs expressed *CCL2*, *CCL7* and *IL6*. However, because the BAL procedure does not sample epithelial, endothelial or stromal cells, we cannot interpret these cells as the primary source of these cytokines. For example, eotaxin-1 (*CCL11*) – an important chemokine involved in eosinophil recruitment – was not expressed in any of the cells sampled by the BAL procedure, but it is known to be expressed by peribronchial and alveolar fibroblasts^25^, suggesting activation of these cells as a result of unresolved lung injury.

### Microbiologic analysis of BAL fluid reveals a respiratory pathogen in a significant minority of patients

All BAL samples collected in this cohort were sent to the clinical laboratory for analysis with multiplex PCR, quantitative culture and other testing as clinically indicated (**Supplemental Table 4**). We detected SARS-CoV-2 virus in the BAL fluid of six patients with RPRA and in three of these patients, SARS-CoV-2 transcripts were also detected using single-cell RNA-seq (**Supplemental Figure 4f**). Three of these patients were on immunosuppression after solid organ or stem cell transplant, one was on ibrutinib for chronic lymphocytic leukemia (CLL), one was on rituximab for rheumatoid arthritis, and one had multiple medical comorbidities including heart failure and in remission for prostate cancer and lymphoma. One of the patients in whom SARS-CoV-2 was detected by both single-cell RNA-seq and PCR received paxlovid accompanied by a dose reduction in prednisone with complete recovery of respiratory symptoms and improvement in CT imaging of the chest, and one died.

Two patients had a respiratory bacterial pathogen identified by multiplexed PCR and confirmed by quantitative culture (*P. aeruginosa* and *S. aureus*). One sample was PCR positive for Human rhinovirus/enterovirus on multiplex PCR, this sample was also positive for SARS-CoV-2 by PCR. Quantitative culture of BAL fluid identified respiratory pathogens not typically seen as part of oral flora in six patients, two of these samples were also positive for SARS-CoV-2. Microbiologic results for the cohort can be found in **Supplemental Table 4**.

### Transcriptomic changes in the nasal mucosa do not reflect unresolved lung injury in the distal lung in patients with RPRA

Several studies have demonstrated that transcriptomic changes in airway epithelium, including epithelium in nasopharynx, can reflect disease-associated processes in the distal lung parenchyma. Therefore, in a subset of 5 patients with RPRA and 6 healthy volunteers, we obtained nasal curettage samples at the time of bronchoscopy, which were processed and analyzed using single-cell RNA-seq. We resolved expected epithelial populations as well as several immune cell populations (**Figure 6a–c, Supplemental Figure 6a, Supplemental Data Files 14–17**). The abundance of epithelial and immune cell populations did not differ between patients with RPRA and healthy volunteers (**Figure 6d**). Similarly, we found very few differentially expressed genes between the groups and no differentially expressed Spectra programs (**Figure 6e, Supplemental Figure 6b**).

**Figure 6:**
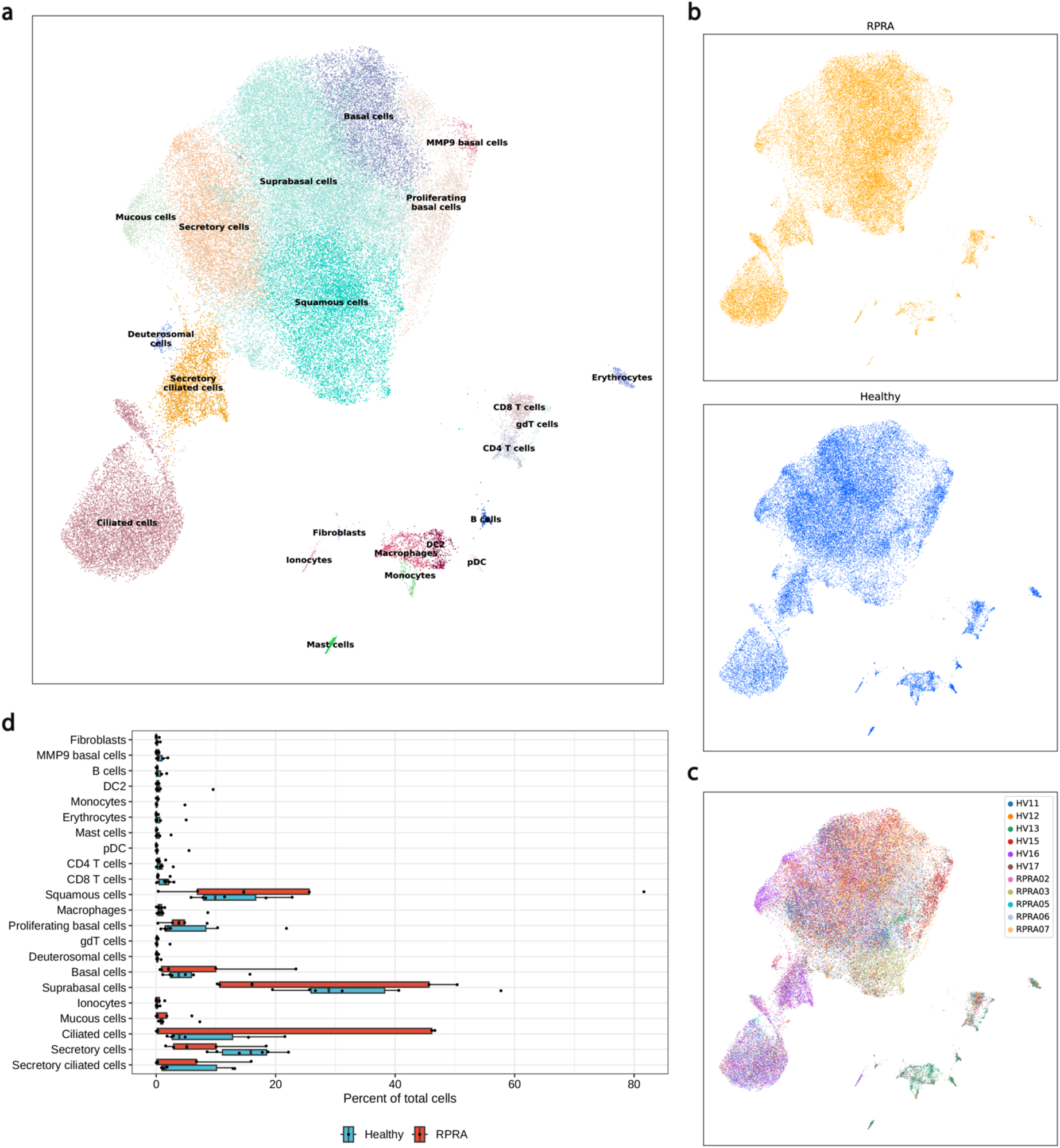
Transcriptomic changes in the nasal mucosa do not reflect ongoing inflammation in the distal lung in patients with RPRA. **a.** UMAP plot showing integrated single-cell RNA-seq analysis of nasal mucosa from 5 patients with RPRA and 6 healthy volunteers, split by diagnosis (**b**) and subject (**c**). **d**. Proportions of different cell types in nasal mucosa between healthy controls and patients with RPRA. No differences are significant (q < 0.05, pairwise Wilcoxon rank-sum tests with FDR correction).

## DISCUSSION

Respiratory symptoms are among the most common complaints in patients who present with PASC but whether the symptoms and signs that define the syndrome of PASC result from a common pathobiology or represent a collection of distinct molecular disorders with shared symptoms is not known. Here we focused on a group of patients with relatively severe manifestations of PASC characterized by persistent respiratory symptoms and detectable abnormalities on computed tomography chest imaging. The patients in this cohort showed substantial heterogeneity in their clinical course after diagnosis with SARS-CoV-2 pneumonia and their clinical and radiographic presentation. Despite this heterogeneity, analysis of bronchoalveolar lavage fluid (BAL) revealed shared molecular features that suggest a common pathobiology. Using a systems approach that integrated data from flow cytometry, BAL fluid cytokines and single-cell RNA-seq of BAL fluid samples, we found the severity of radiographic abnormalities was associated with persistent inflammatory response in the alveolar space characterized by alveolar neutrophilia and the ongoing recruitment of profibrotic monocyte-derived alveolar macrophages (MoAM). In some patients, persistent infection with SARS-CoV-2 or secondary bacterial infection, readily detected by analysis of BAL fluid, were possible drivers of the ongoing inflammation.

A machine learning classifier characterized most of the CT abnormalities in our cohort as more likely suggestive of lung fibrosis than lung inflammation and the improvement on follow up CT scans we observed in most patients was primarily attributable to reduced fibrosis. This is reminiscent of animal models of fibrosis induced by bleomycin or viral infection, which also resolve spontaneously. Using the bleomycin model, we and others have identified a multicellular model of fibrosis that begins with injury to the alveolar epithelium^12, 16^, as also occurs with SARS-CoV-2 infection^39^. This results in the recruitment of classical monocytes from the circulation to the alveolar space where they differentiate into profibrotic MoAM^9, 10, 12–14^. In murine models of lung injury and fibrosis, we and others have shown deletion of profibrotic MoAM ameliorates disease severity, causally implicating them in the pathobiology of fibrosis^10, 12, 13^. Furthermore, pharmacologic strategies that accelerate alveolar epithelial repair reduce the recruitment and accelerate the maturation of profibrotic MoAM^16^. These animal data suggest the profibrotic MoAM we detected in patients with RPRA might serve as both biomarkers of failed alveolar epithelial repair and as targets for future therapy.

We identified persistent SARS-CoV-2 infection as detected by PCR analysis of BAL fluid samples and single-cell RNA-seq analysis in 6 patients, all of whom were immunocompromised. In addition, clinical analysis of samples from bronchoscopy specimens identified bacterial co-infections in some patients. These data highlight the potential utility of bronchoscopy prior to the initiation of empiric therapy in patients with PASC to exclude infection. One patient with persistent SARS-CoV-2 was treated with paxlovid combined with reduced immune suppression associated with improvement. Further study is needed to determine whether and how to administer antiviral therapies for these patients.

To date, only a handful of studies have performed single cell analysis of alveolar space in patients with respiratory PASC and attempted to link it to objective radiographic changes^40, 41^. While these studies identified changes in T cells in BAL fluid from patients with respiratory PASC, they were unable to link specific immunological abnormalities to fibrotic pathology. In contrast, our finding of that abundance of profibrotic MoAM correlates with the extent of fibrotic abnormalities in patients with RPRA provides important insights into its pathobiology. Our study was underpowered, however, to evaluate the abundance of these cells, their transcriptional signatures, the co-occurrence of other immune cells (e.g., Tregs), or levels of BAL fluid cytokines as biomarkers. Nevertheless, our data can be used to inform the development of lower cost flow cytometry or limited PCR panels that can be applied to bronchoscopy specimens. Unfortunately, although sampling by nasal curettage offers a less invasive method to longitudinally assess clinical responses to therapy, our single cell analysis of nasal curettage samples suggests the resolution of inflammation in the nasal epithelium is largely independent from the alveolar compartment. Although our inability to detect transcriptomic changes in the nasal mucosa in patients with RPRA might be due to a small sample size.

Our study has several limitations. First, because this was an observational cohort of limited size, we cannot make conclusions about the impact of patient-specific comorbidities or COVID-19-specific therapies that might have contributed to the course of RPRA. Second, because almost all patients in the cohort received corticosteroids, we cannot determine whether their administration, or other therapies the patients received, affected the improvement we observed in the majority of the cohort. Third, our study is insufficient by itself to recommend bronchoscopic sampling to guide therapy for patients with RPRA, but it should contribute to equipoise around this question. Finally, the duration of illness in COVID-19 exceeds that associated with other respiratory viral infections^9, 42^. Hence, while persistent respiratory symptoms and CT abnormalities have long been observed in survivors of pneumonia caused by influenza and other respiratory viruses^43^ and we cannot say whether the pathobiology in these patients is similar to survivors of COVID-19. This problem will be challenging to study as only a handful of these patients are seen at any center each year. Integrating our single-cell RNA-seq data from BAL fluid and nasal curettage samples from patients with PASC with those from patients with persistent symptoms after pneumonia secondary to other respiratory viruses, or new strains of SARS-CoV-2, provides a possible approach to address this question across centers.

## METHODS

### Human subjects

All human subjects research was approved by the Northwestern University Institutional Review Board. Patients with RPRA were enrolled in the study STU00213592. Healthy volunteers were enrolled in the study STU00206783 and STU00214826 at Northwestern University, or Pro00088966 and Pro00100375 at Duke University. Two patients with severe lung fibrosis necessitating consideration for lung transplant after COVID-19 were co-enrolled in STU00212120 and STU00213592. All study participants or their surrogates provided informed consent.

### Patient characteristics

35 patients were enrolled after undergoing evaluation at Northwestern Memorial Hospital between November 2020 and April 2022. Two patients were evaluated as inpatients; the remaining 33 patients were seen in the outpatient setting for symptoms related to RPRA. All patients enrolled in this study had a history of acute COVID-19 infection (nasopharyngeal swab PCR positive), persistent respiratory symptoms, and abnormal CT lung imaging at least three months after COVID-19 diagnosis. Two patients in the cohort subsequently underwent lung transplantation for COVID-19-induced lung fibrosis.

Patients underwent chest radiography, pulmonary function testing, laboratory assessment, and in person or telehealth visits at the discretion of the treating physician. Bronchoscopy was usually performed to exclude ongoing COVID-19 infection or superimposed respiratory infections as a cause of persistent pulmonary symptoms and radiographic abnormalities prior to initiation of glucocorticoids. Clinical data was collected by chart review.

### Bronchoscopy and bronchoalveolar lavage (BAL)

Bronchoscopic BAL was performed in patients in the bronchoscopy suite or in the intensive care unit. Patients were given sedation and topical anesthesia at the discretion of the physician performing the bronchoscopy. The most involved bronchopulmonary segment was identified based on clinician based on review of the chest CT scan and 90–120 ml of saline was instilled into the segment of interest and aspirated back with the first 5 cc of return discarded. Residual fluid beyond that needed for clinical testing (cell count, differential, BioFireTM Pneumonia Panel multiplex PCR, amylase and quantitative culture) was refrigerated on site and processed in less than six hours. The BAL sample from the donor lung in patient RPRA13 was included in the analysis together with BAL samples obtained from healthy volunteers.

### Nasal curettage

Briefly, donors were seated and asked to extend their neck. A nasal curette (Rhino-Pro; VWR) was inserted into either naris, and gently slid in the direction of posterior to anterior ∼1 cm along the lateral inferior turbinate. Five curettes were obtained per participant. The curette tip was then cut and placed in 2 ml hypothermosol and stored at 4°C until processing.

### Treatment and follow-up assessment

The standard corticosteroid regimen was 1 mg/kg of prednisone, tapered by 10 mg every 2 weeks. For all patients undergoing steroid treatment, a follow up CT of the chest was available.

### CT scan machine learning analysis

We evaluated each baseline high-resolution computed tomography (HRCT) scan using previously established quantitative techniques^18, 19^ using the Chest Imaging Platform (www.chestimagingplatform.org). In brief, after segmenting the lungs and the lobes, the lung parenchyma of every scan was analyzed by classifying regions of interest into one of three categories: normal lung, interstitial alterations, or emphysema. This classification was achieved through the utilization of a k-nearest neighbors classifier, which relied on local tissue density and distance from the pleural surface. Parenchymal changes were further categorized into reticular, subpleural line, linear scar, and honeycombing (these aggregate interstitial features hereafter referred to as “fibrotic”); centrilobular nodule and nodular; and ground-glass patterns using the characteristics of the local histogram computed in the patch size of 32×32 pixels. Normal appearing lung parenchyma was further reclassified into high attenuation normal lung for those patches whose mean lung density was above the 95th percentile of lung density from a training subset of control lifelong non-smokers as previously described^44^. Ground-glass patterns and high attenuation normal lung (referred to as “normal inflamed”) were aggregated into the inflammatory compartment. The total lung volume was used to standardize each feature, which was then aggregated for analysis.

### Flow cytometry and cell sorting

BAL fluid samples were filtered through a 70 μm cell strainer, pelleted by centrifugation at 400 rcf for 10 min at 4°C, followed by hypotonic lysis of red blood cells with 2 ml of PharmLyse (BD Biosciences) reagent for 2 minutes. Lysis was stopped by adding 13 ml of MACS buffer (Miltenyi Biotech). Cells were pelleted again and resuspended in 100 μl of 1:10 dilution of Human TruStain FcX (Biolegend) in MACS buffer, and a 10 μl aliquot was taken for counting using K2 Cellometer (Nexcelom) with AO/PI reagent.

The cell suspension volume was adjusted so the concentration of cells was always less than 5×10^7^ cells/ml and the fluorophore-conjugated antibody cocktail was added in 1:1 ratio (**Supplemental Table 5**). After incubation at 4°C for 30 minutes, cells were washed with 5 ml of MACS buffer, pelleted by centrifugation, and resuspended in 500 μl of MACS buffer with 2 μl of SYTOX Green viability dye (ThermoFisher). Cells were sorted on a FACS Aria III SORP instrument using a 100-μm nozzle at 20 psi. Cells were sorted into 300 μl of 2% BSA in DPBS and cryopreserved using the protocol by Linas Mazutis^45^. Briefly, cells pelleted by centrifugation at 400 rcf for 5 min at 4°C, resuspended in Bambanker freezing media to ∼2000 cells/μl concentration. Concentration was confirmed using K2 Cellometer (Nexcelom) with AO/PI reagent using “Immune cells low RBC” program with default settings and ∼40 μl aliquots were immediately frozen at -80°C. Sample processing was performed in BSL-2 facility using BSL-3 practices. Analysis of the flow cytometry data was performed using FlowJo 10.6.2. using uniform sequential gating strategy reported in our previous publication^9^ and reviewed by two investigators (SS, AVM). Relative cell type abundance was calculated as a percent out of all singlets/live/CD45+ cells.

### Single-cell RNA-seq of flow cytometry-sorted BAL cells

Single-cell RNA-seq was performed using Chromium Next GEM Single Cell 5’ reagents V2 (10x Genomics, protocol CG000331 Rev A). Immediately before loading 10x Genomics Chip K with Chromium Single Cell 5’ gel beads and reagents aliquots of cryopreserved cells were retrieved from -80°C freezer, rapidly thawed in water bath at 37°C, gently mixed by pipetting and added to RT mix. The volume of the single cell suspension was calculated using protocol CG000331 Rev A (10x Genomics) based on concentration at the time of cryopreservation and aiming to capture 5000–10000 cells per library. Libraries were prepared according to the manufacturer’s protocol (10x Genomics, CG000331 Rev A). After quality checks, single-cell RNA-seq libraries were pooled and sequenced on a NovaSeq 6000 instrument.

### Nasal curettage processing and single-cell RNA-seq

A single-cell suspension was generated using the cold-active dispase protocol reported by Deprez et al.^46^ and Zaragosi and Barbry^47^ with slight modification. Specifically, ethylenediaminetetraacetic acid (EDTA) was omitted and cells were dispersed by pipetting 20 times every 5 min using a 1 ml tip instead of tritration using a 21/23 G needle. The final concentration of protease from *Bacillus licheniformis* was 10 mg/ml. The total digestion time was 30 min. Following the wash in 4 ml 0.5% BSA in PBS and centrifugation at 400 g for 10 min, cells were resuspended in 0.5% BSA in PBS and counted using a Nexcelom K2 Cellometer with acridine orange/propidium iodide reagent. This protocol typically yields ∼300– 500,000 cells with a viability of >95%. The resulting single-cell suspension was then used to generate single-cell libraries following the protocol for 5′ V1 (CG000086 Rev M; 10x Genomics) or V2 chemistry (CG000331 Rev A; 10x Genomics). Excess cells from two of the samples from healthy volunteers were pooled together to generate one additional single-cell library. After a quality check, the libraries were pooled and sequenced on a NovaSeq 6000 instrument. To assign sample information to cells in the single-cell library prepared from two samples, we ran souporcell version 2.0^48^ for that library and two libraries that were prepared from these samples separately. We used common genetic variants prepared by the souporcell authors to separate cells into two groups by genotype for each library, and Pearson correlation between the identified genotypes across libraries to establish correspondence between genotype and sample. We prepared, sequenced, and analyzed single-cell RNA-seq libraries from seven healthy volunteers. However, after initial analysis one library was excluded due to overall low quality (HV14), and only six libraries were included in the final analysis.

### Statistical methods

All statistics in the manuscript are reported as mean ± standard deviation. When multiple hypothesis tests were performed, the false discovery rate (FDR) was controlled using the procedure of Benjamini and Hochberg^49^. A significance level of 0.05 was used for all tests. Wilcoxon rank-sum tests and permutation tests were performed in R using coin 1.4-2^50, 51^, and in Python using scipy 1.10.0^52^. Linear models and associated confidence intervals were computed using the stat_smooth function with method = “lm” in ggplot2 3.4.2^53^.

### Single-cell RNA-seq analysis

Data was processed using the Cell Ranger 7.0.0 pipeline with exon-only processing mode (10x Genomics). To enable detection of viral RNA, reads were aligned to a custom hybrid genome containing GRCh38.98 and SARS-CoV-2 (NC_045512.2). An additional negative-strand transcript spanning the entirety of the SARS-CoV-2 genome was then added to the GTF and GFF files to enable detection of SARS-CoV-2 replication. Data were processed using Scanpy 1.9.2^54^, and multisample integration was performed with scvi-tools 0.20.0^55–57^. The scVI models for both the BAL and nasal samples were constructed on 1000 HVGs with the hyperparameters n_layers = 2, dropout_rate = 0.2, and n_latent = 10, and were trained using the settings max_epochs = 500, check_val_every_n_epoch = 2, and early_stopping = True. Default hyperparameters and settings were used otherwise. An initial round of Leiden clustering using the function sc.tl.leiden was performed on the integrated BAL object with a resolution of 1.2, and on the integrated nasal object with a resolution of 0.75. Clusters characterized by low number of detected genes and transcripts and high percentage of mitochondrial genes were removed. Clusters containing doublets were identified as clusters simultaneously expressing lineage-specific marker genes (for example *C1QA* for macrophages and *CD3G* for T cells) and excluded. Cell types were identified by marker genes, computed using the sc.tl.rank_genes_groups function with the settings method = “wilcoxon”, n_genes = 200, and default settings otherwise. Count matrices are available via GEO: BAL from healthy volunteers GSE232616, nasal curettage from healthy volunteers GSE232623, and BAL and nasal curettage from patients with RPRA GSE232627.

### Differential abundance analysis for single-cell RNA-seq data

Since neutrophils represented a significant fraction of immune cells in BAL fluid and were excluded during the cell sorting, we used the percentage of neutrophils from flow cytometry analysis to correct the denominator for estimating relative cell abundance. See code for details.

### Differential expression analysis

To take advantage of the multiple subjects in each condition and avoid p-value inflation inherent to approaches where each cell is treated as an independent observation, we summed RNA transcript counts for each subject on per cell type level (pseudobulk approach). Samples were retained for differential expression analysis of a given cell type if they contained at least 40 cells of that type, and if they constituted at least 1% of all cells of that type. Differential expression analysis was performed in R 4.1.1 using DESeq2 1.34.0. A ‘local’ model of gene dispersion was used; default settings were used otherwise. Differentially expressed genes were those with q < 0.05 (Wald test with FDR correction). To help identify genes of interest, we applied two filtering criteria. First, we removed genes encoding ribosomal proteins the list of differentially expressed genes. The second criterion was used to correct for the fact that information on number of cells is not taken into account in pseudobulk differential expression. For each gene and cell type, we counted the number of samples in which the gene was expressed in more than 2% of cells of that type in each sample. Genes which did not satisfy this criterion in at least four samples were removed.

### Factor analysis of single-cell RNA-seq data

Matrix factorization was performed using Spectra 0.1.0^34^. The list of gene sets used for the initial factors is the same as the list used in the Spectra manuscript^34^, and are provided here (**Supplemental Data File 8**). Factors which had less than three genes expressed in the data were removed. Training for both the BAL and nasal data was done using the hyperparameters lam = 0.01, rho = None, and default hyperparameters and settings otherwise. To determine which factors were differentially expressed between RPRA subjects and healthy volunteers in each cell type, we defined a subject score as the mean cell score of a factor across all cells of the specified type belonging to each subject. Differentially expressed factors were those with q < 0.05 (Wilcoxon rank-sum test on subject scores with FDR correction). Samples were used in differential factor expression analysis for a given cell type if they contained at least 40 cells of that type, and if they constituted at least 1% of all cells of that type. A minimum of three subjects per condition were required for comparison. When computing correlations between subject scores and CT features, a minimum of ten RPRA subjects were required to pass the filtering procedure to ensure robust correlations.

### Multiplexed cytokine assays of BAL fluid

Multiplexed cytokine profiling of BAL fluid was performed by Eve Technologies (Calgary, Alberta, Canada). Samples were thawed and aliquoted at 100 µL, frozen and shipped to the CRO on dry ice. The Human Cytokine/Chemokine 71-Plex Discovery Assay (HD71) was then performed on each sample. Custom outputs containing raw median fluorescent intensity (MFI) values, standard curve concentrations, and bead counts for processing as described below.

### Multiplexed cytokine assay processing and analysis

Analysis was performed as described in our previous publication^58^. Briefly, raw MFI values, beads counts, and standard concentrations were first stripped from the data output from either Exponent (in-house assays; Luminex) or bespoke output from Eve Technologies (Calgary, Alberta, Canada). MFI measurements with fewer than 50 bead counts were discarded. Standard curves for each cytokine were then fit for each assay run using self-starting 5-parameter logistic (5PL) models using drc 3.2-0^59^. Cutoffs for curves with low predictive value were then determined empirically using histograms MFI values vs standard concentrations to identify a bimodal distribution cutoff. For in-house assays, all values calculated using standard curves with MFI < 50 at 100 pg/mL were discarded. For Eve Technologies assays, all values calculated using standard curves with MFI < 50 at 10 pg/mL were discarded. Experimental values for each cytokine were then predicted using the ED function in drc with “absolute” value prediction. In rare cases where a 5PL model could not be fit for an individual cytokine-assay combination, these values were excluded. Values below the lower asymptote of the model were set to a concentration of 0 pg/mL. Values above the upper asymptote were set to the value of the upper asymptote. Technical replicates were collapsed by mean with NA values excluded.

## Code availability

Code is available at https://github.com/NUPulmonary/Bailey_Puritz_RPRA_2023. Single-cell RNA-seq data can be explored via the UCSC Cell Browser at https://www.nupulmonary.org/covid-19/.

## ACKNOWLEDGEMENTS

This research was supported in part through the computational resources and staff contributions provided for the Quest high performance computing facility at Northwestern University which is jointly supported by the Office of the Provost, the Office for Research, and Northwestern University Information Technology.

This research was supported in part through the computational resources and staff contributions provided by the Genomics Compute Cluster which is jointly supported by the Feinberg School of Medicine, the Center for Genetic Medicine, and Feinberg’s Department of Biochemistry and Molecular Genetics, the Office of the Provost, the Office for Research, and Northwestern Information Technology. The Genomics Compute Cluster is part of Quest, Northwestern University’s high performance computing facility, with the purpose to advance research in genomics.

Northwestern University Flow Cytometry Core Facility is supported by NCI Cancer Center Support Grant P30 CA060553 awarded to the Robert H. Lurie Comprehensive Cancer Center. Cell sorting was performed on a BD FACSAria SORP cell sorter purchased through the support of NIH 1S10OD011996-01.

Integrative genomic services were performed by the Metabolomics Core Facility at Robert H. Lurie Comprehensive Cancer Center of Northwestern University.

Next generation sequencing was performed with support from Simpsons Querrey Institute for Epigenetics. This work was supported, in part, by a generous gift from Mr. and Mrs. Michael Ferro.

J.I.B. was supported by NIH grants T32HL076139 and UL1TR001422. C.H.P and R.B. were supported by NIH/NIA R01AG068579, Simons Foundation 597491-RWC01, and NSF 1764421-01. R. San Jose Estepar was supported by NIH 5R21LM013670. A.B. was supported by NIH grants R01HL147290, R01HL145478, and R01HL147575. R.A.G. was supported by NIH grant 1F31AG071225. R.M.T. was supported by NIH grants R01ES034350 and R01ES027574. K.M.G. was supported by NIH grant R01ES028829. K.M.R. was supported by NIH grants P01HL154998 and P01AG049665. G.R.S.B was supported by a Chicago Biomedical Consortium grant, Northwestern University Dixon Translational Science Award, Simpson Querrey Lung Institute for Translational Science (SQLIFTS), and NIH grants AG049665, HL154998, HL14575, HL158139, HL147290, AG075423, AI135964 and The Veterans Administration award I01CX001777. A.V.M was supported by NIH grants U19AI135964, P01AG049665, P01HL154998, R01HL153312, R01HL158139, R01ES034350, R21AG075423.

---

**Supplement Figure 1:**
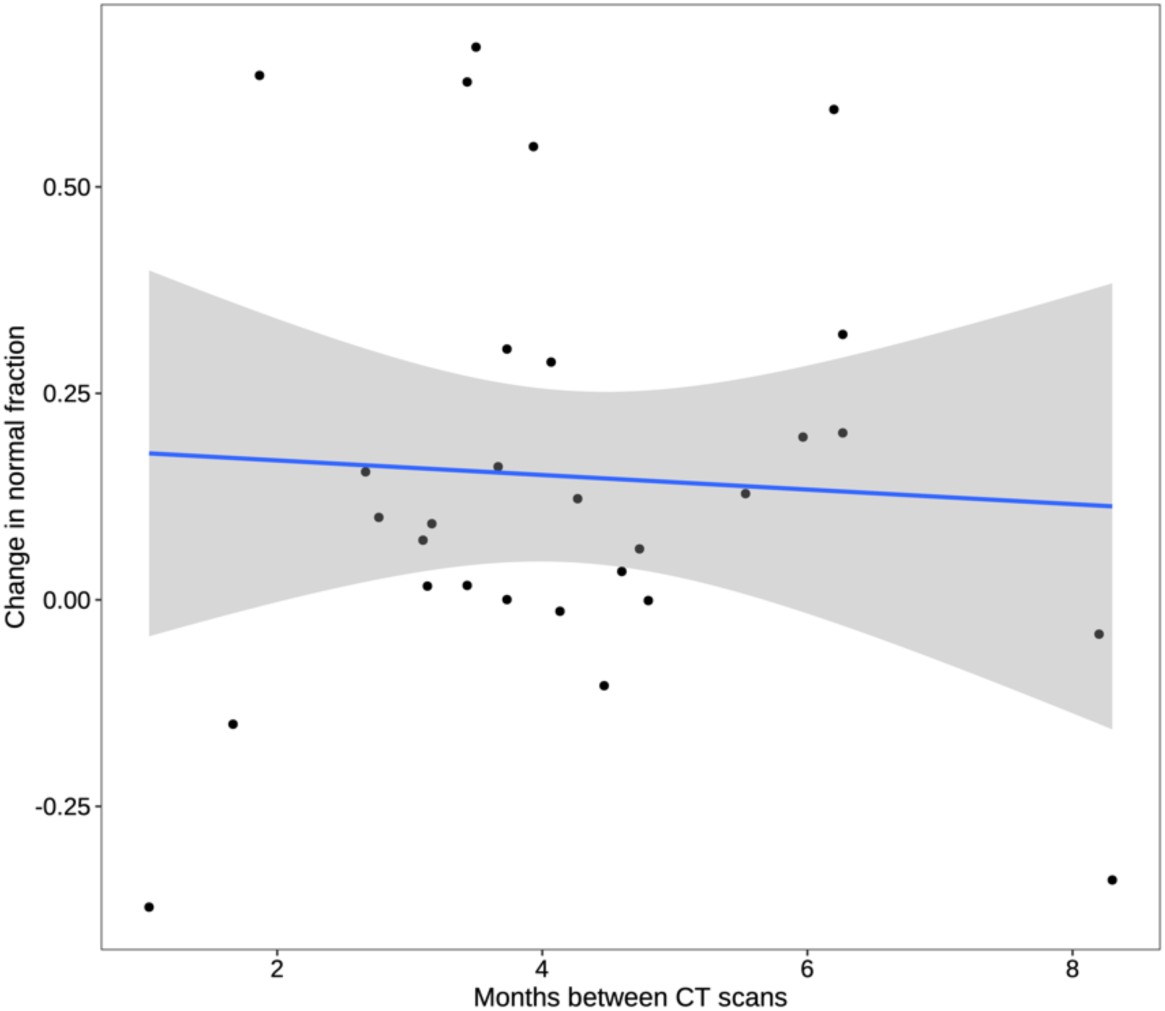
Improvement of abnormalities on CT imaging does not correlate with time. Comparison of the change in normal lung fraction with the interval between the initial and follow-up CT scans. The correlation (Spearman’s rho) is small (ρ = 0.01) and is not significant (permutation test, p = 0.96). A linear model and 95% confidence interval are shown.

**Supplemental Figure 2:**
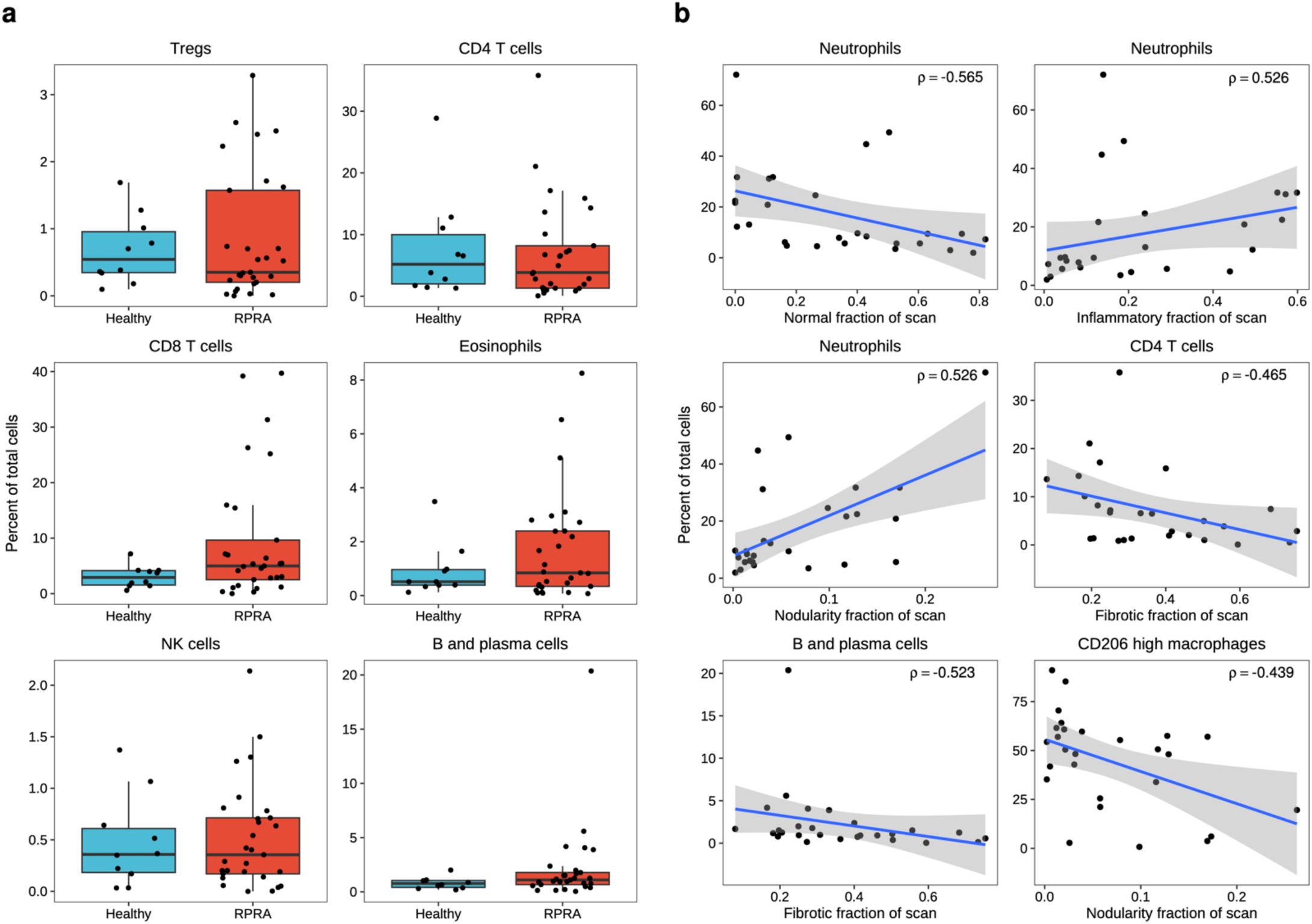
Comparison of cell populations detected using flow cytometry analysis of BAL fluid from patients with RPRA and healthy controls. **a.** Proportions of cells in BAL fluid that were not differentially abundant between healthy controls and patients with RPRA (including patients who required lung transplant) (q < 0.05, pairwise Wilcoxon rank-sum tests with FDR correction). **b.** Comparison of cell abundances for the described cell populations with the described CT abnormalities. Only associations with |ρ| > 0.4 are shown. No associations were statistically significant (q < 0.05, permutation tests with FDR correction). Linear models and 95% confidence intervals are shown.

**Supplemental Figure 3:**
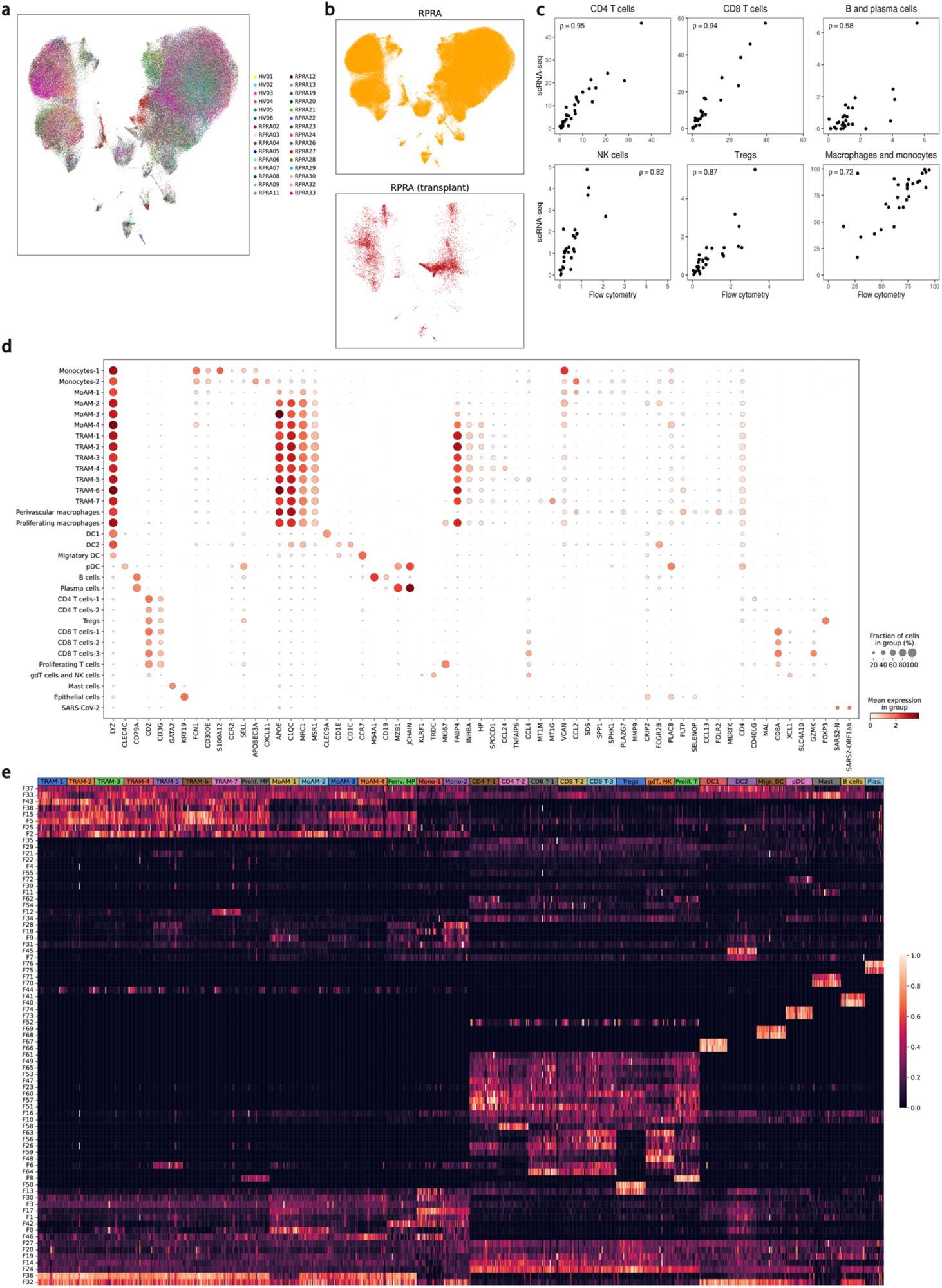
Ongoing recruitment of profibrotic MoAM is associated with fibrotic abnormalities on CT scans. **a–b.** UMAP plot showing integrated analysis of BAL immune cells from 24 patients with RPRA and 6 healthy volunteers, split by subject (**a**), and transplant status (**b**). **c.** Correlation (Spearman’s rho) between cell type abundances determined by flow cytometry and single-cell RNA-seq. Correlation coefficients are annotated on each plot. **d.** Dot plot showing expression of the key cell type markers used to identify cell types. **e.** Heatmap of subject scores for Spectra programs within each cell type. Each column represents a single subject. Rows are min-max scaled. Prolif., proliferating; MP, macrophage; periv., perivascular; mono, monocyte; migr., migratory; plas., plasma.

**Supplemental Figure 4:**
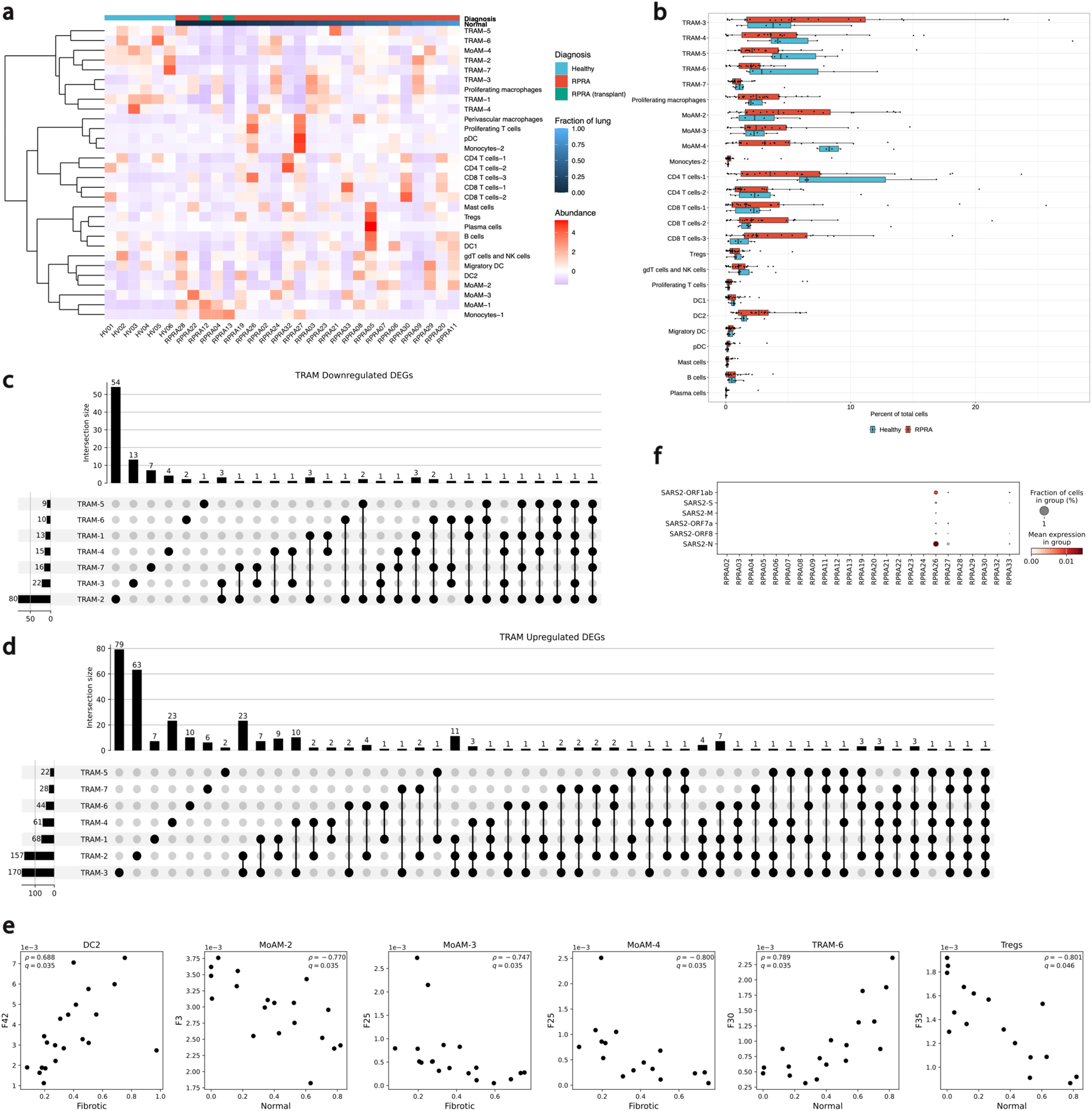
Differential gene expression in BAL macrophages is associated with fibrotic abnormalities on CT scans. **a.** Hierarchical clustering of cell type abundances from BAL samples from healthy volunteers and patients with RPRA. Column headers are color-coded by the diagnosis. Samples from the two patients with RPRA who required transplant are coded separately. RPRA subjects are sorted by the fraction of normal lung on the initial CT scan. Clustering was performed using Ward’s method. Rows are z-scored. **b.** Proportions of cells in BAL fluid that were not differentially abundant between healthy controls and patients with RPRA (including patients who required lung transplant) (q < 0.05, pairwise Wilcoxon rank-sum tests with FDR correction). **c.** Upset plot showing downregulated genes shared between TRAM subsets. **d.** Upset plot showing upregulated genes shared between TRAM subsets. **e.** Correlations (Spearman’s rho) between Spectra subject scores and CT features in specified cell populations. Only significant associations are shown (q < 0.05, permutation tests with FDR correction). Adjusted p-values and correlation coefficients are annotated on each plot. **f.** Dot plot showing expression of SARS-CoV-2 genes.

**Supplement Figure 5:**
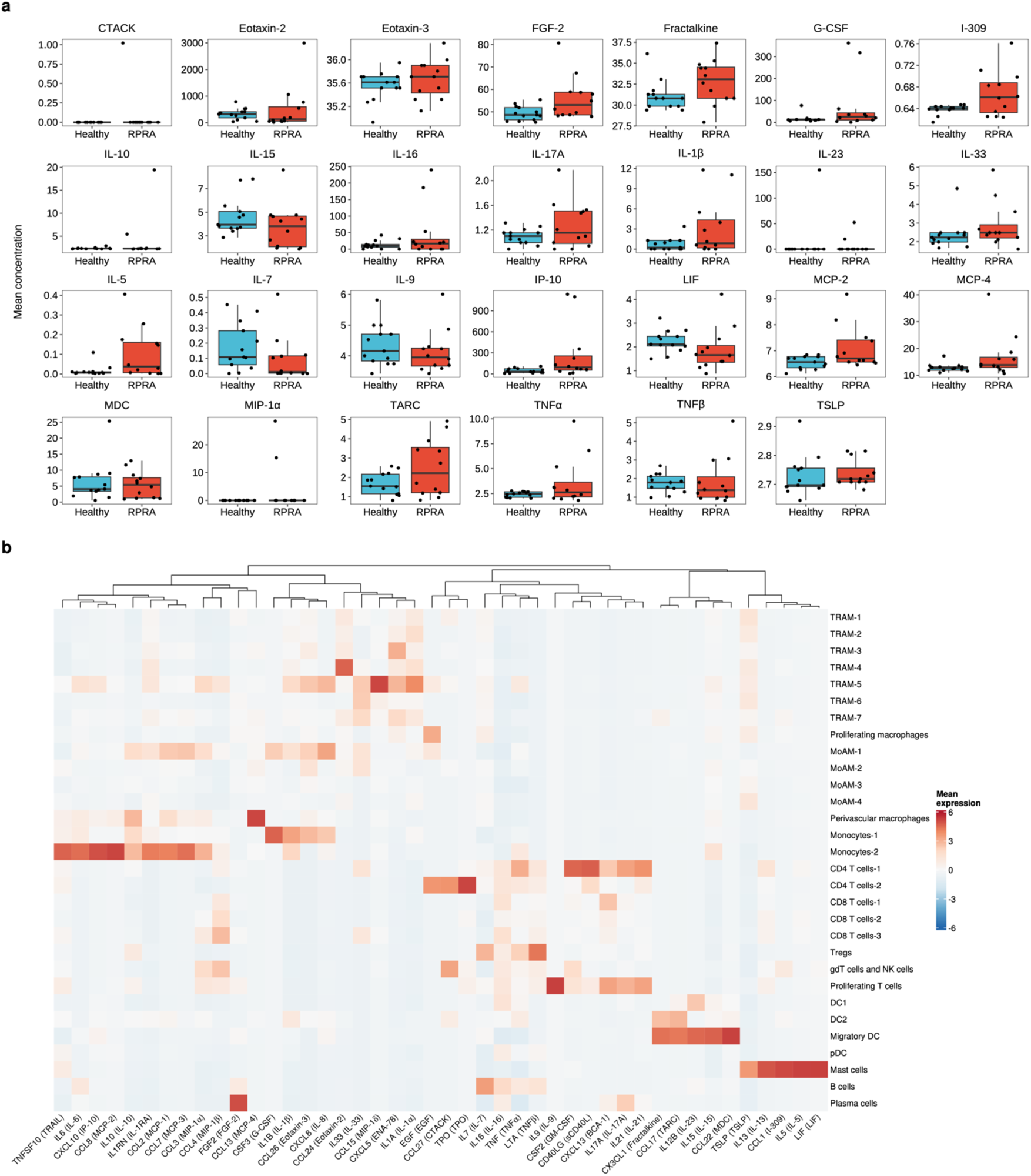
**a.** Levels of cytokines or chemokines that did not differ significantly (q < 0.05, pairwise Wilcoxon rank-sum tests with FDR correction) between patients with RPRA (including patients who required lung transplant) and healthy controls. **b.** Hierarchical clustering of mean expression levels from BAL single-cell RNA-seq data of genes encoding each cytokine measured. *CCL11* (eotaxin-1), *IL20* (IL-20), and *CCL21* (6CKine) are not shown as these genes were not expressed in cells sampled via BAL procedure. Clustering was performed using Ward’s method. Columns are z-scored.

**Supplemental Figure 6:**
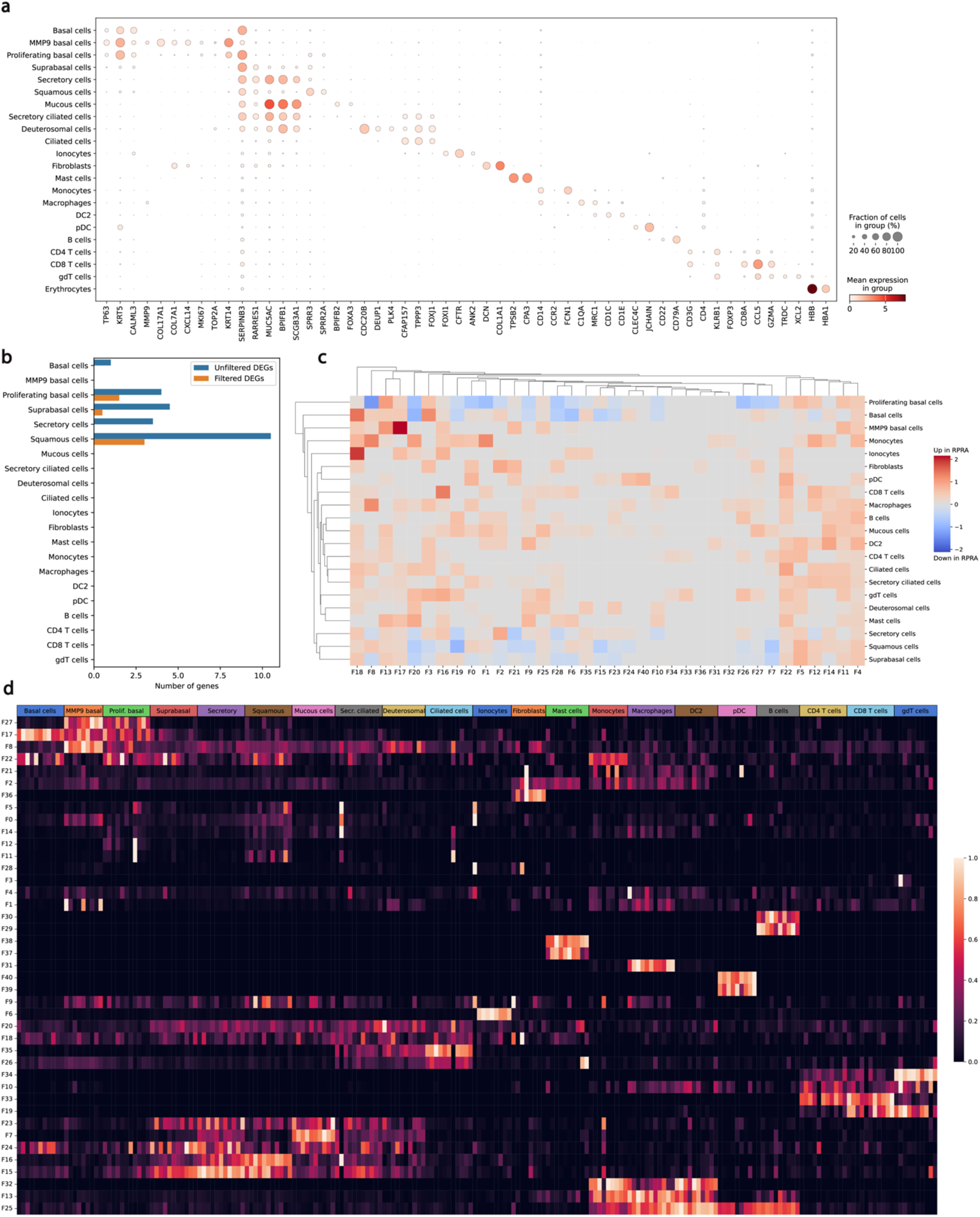
Transcriptomic changes in the nasal mucosa do not reflect ongoing inflammation in the distal lung in patients with RPRA. **a**. Dot plot showing expression of the key cell type markers used to identify cell types. **b.** Bar plot showing number of differentially expressed genes between different cell types in nasal mucosa in healthy controls and patients with RPRA (q < 0.05, Wald test with FDR correction) with and without filtering criteria applied (see methods). **c.** Hierarchical clustering on the signal-to-noise ratio of Spectra subject scores between patients with RPRA and healthy controls. No factors were differentially expressed (q < 0.05, Wilcoxon rank-sum test on subject scores with FDR correction). **d.** Heatmap of subject scores for Spectra programs. Each column represents a single subject. Rows are min-max scaled.

**Supplemental Table 1.** Demographics of patients in the cohorts. Continuous demographics are reported as median (minimum - maximum). HV, healthy volunteers. The nasal control subject who was omitted from analysis (HV14) is not included.

| Demographics | RPRA | BAL Flow HV | BAL scRNA-seq HV | BAL Cytokine HV | Nasal scRNA-seq HV |
| --- | --- | --- | --- | --- | --- |
| Female sex | 15/35 (42.9%) | 5/8 (62.5%) | 4/6 (66.7%) | 7/13 (53.8%) | 2/6 (33.3%) |
| Age | 62 (54-66.5) | 27.5 (21-68) | 29.5 (20-68) | 25 (21-40) | 34.5 (25-58) |
| BMI | 29.1 (25.4-32.6) | 28.0 (20.6-35.4) | 24.2 (20.6-34.0) | 24.1 (18.4-35.1) | 22.7 (19.2-27.5) |
| Race |  |  |  |  |  |
| Black or African American | 6/35 (17.1%) | 2/8 (25%) | 0/6 (0%) | 1/13 (7.7%) | 1/6 (16.7%) |
| Asian | 2/35 (5.7%) | 1/8 (12.5%) | 0/6 (0%) | 4/13 (30.8%) | 1/6 (16.7%) |
| White | 23/35 (65.7%) | 5/8 (62.5%) | 6/6 (100%) | 8/13 (61.5%) | 3/6 (50%) |
| Other | 4/35 (11.4%) | 0/8 (0%) | 0/6 (0%) | 0/13 (0%) | 0/6 (0%) |
| Prefer not to answer | 0/35 (0%) | 0/8 (0%) | 0/6 (0%) | 0/13 (0%) | 1/6 (16.7%) |
| Ethnicity |  |  |  |  |  |
| Hispanic or Latino | 7/35 (20%) | 1/8 (12.5%) | 1/6 (16.7%) | 2/13 (15.4%) | 1/6 (16.7%) |
| Not Hispanic or Latino | 28/35 (80%) | 7/8 (87.5%) | 5/6 (83.3%) | 11/13 (84.6%) | 5/6 (83.3%) |

**Supplemental Table 2.**
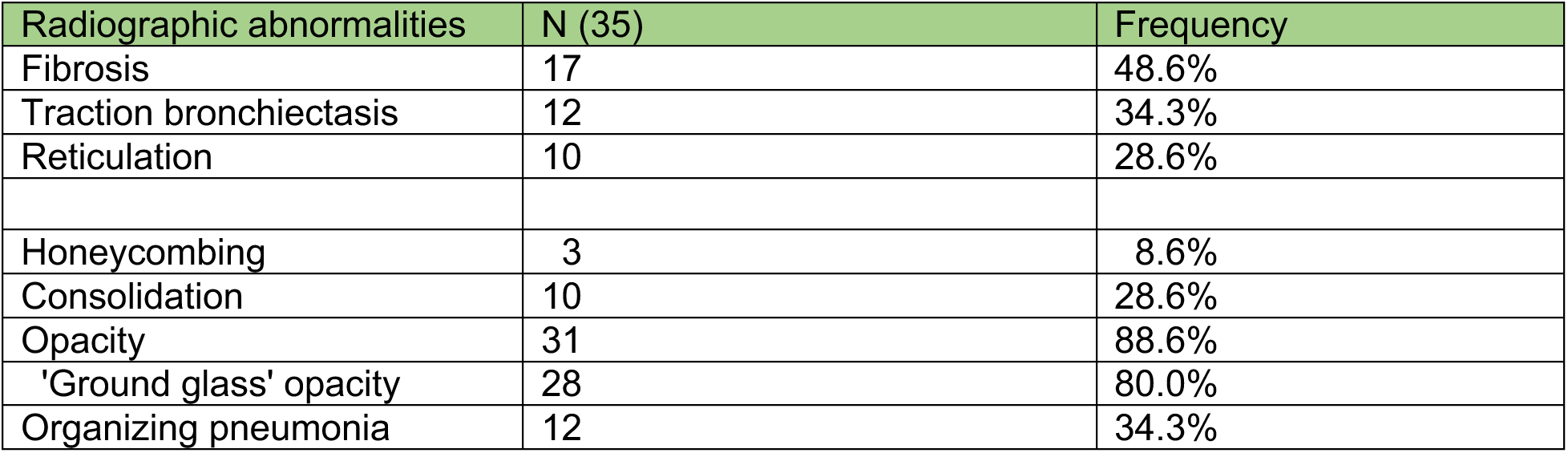
Summary of radiographic features extracted from radiologic reports of initial computed tomography scans (N = 35) of the chest.

| Radiographic abnormalities | N (35) | Frequency |
| --- | --- | --- |
| Fibrosis | 17 | 48.6% |
| Traction bronchiectasis | 12 | 34.3% |
| Reticulation | 10 | 28.6% |
| Honeycombing | 3 | 8.6% |
| Consolidation | 10 | 28.6% |
| Opacity | 31 | 88.6% |
| 'Ground glass' opacity | 28 | 80.0% |
| Organizing pneumonia | 12 | 34.3% |

**Supplemental Table 3.** Grouping of CT abnormalities identified via machine learning analysis.

| Normal | Inflammatory | Fibrotic | Emphysema/Cyst | Nodularity |
| --- | --- | --- | --- | --- |
| Normal not inflamed | Normal inflamed | Honeycombing | Centrilobular emphysema | Nodular |
|  | Ground glass | Linear Scar | Emphysematous | Nodule |
|  |  | Fibronodular | Cyst |  |
|  |  | Reticular |  |  |
|  |  | Subpleural line |  |  |

**Supplemental Table 4.** Results of microbiologic testing by the clinical lab. Two samples were not sent for microbiologic testing. NA indicates the test was not requested by the clinical team. *Streptococcus viridans* 1 and 2 indicate two different strains of *Streptococcus viridans*.

| Study ID | BioFire™ FilmArray Pneumonia Panel | Culture Findings | SARS-CoV-2 PCR | Galactomannan |
| --- | --- | --- | --- | --- |
| RPRA02 | Negative | 15000 CFU *Streptococcus viridans*, 1000 CFU *Enterobacter cloacae*, 3000 CFU *Stenotrophomonas maltophilia*, 100 CFU *Pseudomonas aeruginosa* | Negative | 0.17 |
| RPRA03 | *P. aeruginosa*, negative for carbapenamases and CTX-M | 1000 CFU *Pseudomonas aeruginosa* | Negative | NA |
| RPRA04 | Negative | 30000 CFU *Streptococcus viridans*, 15000 CF *Streptococcus viridans* 2 | Negative | 0.37 |
| RPRA05 | Negative | Negative | Negative | 0.07 |
| RPRA06 | Negative | 15000 CFU *Streptococcus viridans* | Negative |  |
| RPRA07 | *Staphylococcus aureus* | 10,000 CFU *Streptococcus viridans*, 1,000 CFU *Staphylococcus aureus*, 25000 CFU *Neisseria* species | Negative | 0.07 |
| RPRA08 | Negative | 50000 CFU *Streptococcus viridans* 1, 20000 CFU *Streptococcus viridans* 2 | Negative | 0.07 |
| RPRA09 | Negative | 15000 *Streptococcus viridans* | Negative | 0.07 |
| RPRA10 | Negative | Negative | Negative | 0.13 |
| RPRA11 | Negative | 80,000 CFU *Streptococcus viridans* 1, 30000 CFU *Streptococcus viridans* 2, 50000 *Stomatococcus* | Negative |  |
| RPRA12 | NA | 2000 CFU *Staphylococcus aureus* | Negative |  |
| RPRA13 | NA | Negative | Negative | 0.15 |
| RPRA18 | Negative | 10,000 CFU *Streptococcus viridans*, 100 CFU *Klebsiella oxytoca* | Negative | 0.21 |
| RPRA19 | NA | NA | Negative |  |
| RPRA20 | NA | NA | Negative |  |
| RPRA21 | Negative | Negative | Negative |  |
| RPRA22 | Negative | 300 CFU *Enterococcus faecalis* | Negative |  |
| RPRA23 | Human rhinovirus/enterovirus | 40,000 CFU *Streptococcus viridans* | Positive |  |
| RPRA24 | Negative | Negative | Negative | 0.11 |
| RPRA26 | Negative | Negative | Positive |  |
| RPRA27 | Negative | Negative | Positive | 0.07 |
| RPRA28 | Negative | 20000 CFU *Streptococcus viridans* 1, 10000 CFU *Streptococcus viridans* 2 | Negative |  |
| RPRA29 | Negative | Negative | Negative | 0.07 |
| RPRA30 | Negative | 10000 CFU *Streptococcus viridans*, 2000 CFU *Streptococcus pseudopneumoniae* | Positive | 0.1 |
| RPRA32 | Negative | Negative | Negative | 0.08 |
| RPRA33 | Negative | Negative | Positive | 0.06 |
| RPRA34 | Negative | 100 CFU *Klebsiella oxytoca* | Positive | 0.11 |

**Supplemental Table 5.** Flow cytometry panel used for BAL sample phenotyping.

| Antigen | Clone | Fluorochrome | Manufacturer | Cat # |
| --- | --- | --- | --- | --- |
| CD4 | RPA-T4 | BUV395 | BD | 564724 |
| CD19 | HIB19 | BUV395 | BD | 740287 |
| CD25 | 2A3 | BUV737 | BD | 564385 |
| CD56 | NCAM16.2 | BUV737 | BD | 612766 |
| HLA-DR | L243 | eFluor450 | ThermoFisher | 48-9952-42 |
| CD45 | HI30 | BV510 | Biolegend | 304036 |
| CD15 | HI98 | BV786 | BD | 563838 |
| Live/Dead | Not applicable | SYTOX Green | ThermoFisher | S34860 |
| CD3 | SK7 | PE | ThermoFisher | 12-0036-42 |
| CD127 | HIL-7R | PECF594 | BD | 562397 |
| CD206 | 19.2 | PECy7 | ThermoFisher | 25-2069-42 |
| CD8 | SK1 | APC | Biolegend | 344721 |
| CD14 | M5E2 | APC | Biolegend | 301808 |
| EpCAM | 9C4 | APC | Biolegend | 324208 |

**Supplemental Data File 1.** Metadata for RPRA cohort. Dates are reported as days after COVID diagnosis. CSV file.

**Supplemental Data File 2.** Metadata for healthy volunteers. CSV file.

**Supplemental Data File 3.** CT scan data. CSV file.

**Supplemental Data File 4.** Lung fractions from initial CT scans. CSV file.

**Supplemental Data File 5.** Lung fractions from follow-up CT scans. CSV file.

**Supplemental Data File 6.** Cell-type percentages from flow cytometry analysis of BAL fluid. CSV file.

**Supplemental Data File 7.** Marker genes for integrated BAL single cell RNA-seq object. CSV file.

**Supplemental Data File 8.** Input Spectra factors for both BAL and nasal single cell RNA-seq data. XLSX file.

**Supplemental Data File 9.** Gene weights of top 100 genes of Spectra factors for BAL single cell RNA-seq object. CSV file.

**Supplemental Data File 10.** Functional annotation for selected BAL Spectra programs and specific genes. TXT file.

**Supplemental Data File 11.** Cell-type percentages for integrated BAL single cell RNA-seq object. Raw percentages and percentages scaled by the abundance of neutrophils are reported. CSV file.

**Supplemental Data File 12.** Output of DESeq2 analysis for integrated BAL single cell RNA-seq data. CSV file.

**Supplemental Data File 13.** Cytokine concentrations. CSV file.

**Supplemental Data File 14.** Marker genes for integrated nasal single cell RNA-seq object. CSV file.

**Supplemental Data File 15.** Output of DESeq2 analysis for integrated nasal single cell RNA-seq object. CSV file.

**Supplemental Data File 16.** Cell-type percentages for integrated nasal single cell RNA-seq object. CSV file.

**Supplemental Data File 17.** Gene weights of top 100 genes of Spectra factors for nasal single cell RNA-seq object. CSV file.

