## Supplemental Data File 10 for "Expansion of profibrotic monocyte-derived alveolar macrophages in patients with persistent respiratory symptoms and radiographic abnormalities after COVID-19"

Functional annotation of Spectra programs and specific genes.

F43: Genes related to homeostatic alveolar macrophage functions such as lipid metabolism (FABP4, FABP5, APOC2), immune regulation and inflammation (IL1RN, IL1B, IFIT2, IFITM3, ISG15, HLA-DRA, S100A8, TNFAIP3), phagocytosis (MARCO), protease activity (CTSL, CTSW), and antioxidant response (HMOX1, SOD2).

F0: Genes in the program F0 were related to various processes such as cellular signaling and regulation (SGK1, KLF2, KLF6, FOS, FYN), immune response (FCGR2B, HLA-DRA, MS4A6A, TNFSF8, TNFSF14), cell adhesion and cytoskeletal organization (EZR, PHLDA1, LIMD2, CORO1A), extracellular matrix organization and fibrosis (VCAN, SPP1), and protease activity (CTSL, CST3, CFD, PLP2).

F19: Genes in the program F19 were related to cellular signaling and regulation (TSC22D3, KLF6, PLP2, CRIP1, CRIP2, CEBPD, STK17B), cytoskeletal organization, cell cycle, and cell adhesion (EZR, CYTIP, LIMD2, CKLF, STMN1), immune response (NCF1, SKAP1, LCP2, GIMAP7), and RNA regulation (NEAT1, GAS5, ZFP36L1) and DNA and chromatin regulation (KMT2A, SMCHD1).

F42: Genes in the program F42 were related to immune response and inflammation (CCL2, CD14, FCN1, S100A9, CEBPD, S100A12, IFITM3, FCGR2B, NEAT1, CST3, APOBEC3A, S100A8, SGK1, TIMP1, CXCL8), extracellular matrix organization (FOLR2, VCAN), protease activity (LGMN, CTSL, CFD, MMP9, MMP7), cell signaling and transcriptional regulation (MARCKS, FOS, FYB1, TMEM176B, TMEM176A, BATF, KLF2, IFI30, KLF6, MT-ND4, LCP2, TCF4), and iron metabolism (PLTP, HMOX1, SLC40A1).

F35: Wnt signaling pathway (TCF7, LEF1), immune response (TNFRSF4, CORO1A, LCP2, RAC2, BTG1, GAS5, STK17B, IFITM1, NCF1, FYB1), cell adhesion and migration (LIMD2, CD48, SELL, ICAM3), signaling pathways (JAK3, LTB, MARCKSL1, IL32, RASAL3, ISG20).

F30: Genes in program F30 were related to inflammation, immune response, and pathogen clearance (S100A8, S100A9, MS4A6A, NCF1, CKLF, LCP2, MARCO), metabolic regulation (FBP1, RETN, AOC1, SLC18A2, MAOB, PNRC1, HMGB2, PARP8), complement regulation (SERPING1), protein degradation (CTSL), cell cycle (STMN1), cell adhesion and signaling (RAC2, CD48, RASAL3, SKAP1).

F25: Genes in this program were related to immune response and antiviral defense (IFI27, IFIT2, GBP5, NCF1, C1QC), complement regulation (SERPING1), and lysosomal activity and protein degradation (CTSL, CTSW). Together, these findings suggest that diverse subsets of alveolar macrophages contribute to both development and resolution of inflammation and fibrosis.
